# Coalescence-Driven Local Crowding Promotes Liquid-to-Solid-Like Phase Transition in a Homogeneous and Heterogeneous Droplet Assembly

**DOI:** 10.1101/2024.10.08.617323

**Authors:** Chinmaya Kumar Patel, Abhradip Mallik, Deb Kumar Rath, Rajesh Kumar, Tushar Kanti Mukherjee

## Abstract

Liquid-to-solid-like phase transition (LSPT) of disordered proteins *via* metastable liquid-like droplets is a well-documented phenomenon in biology and linked to many pathological conditions including neurodegenerative diseases. However, very less is known about the early microscopic events and transient intermediates involved in the irreversible protein aggregation of functional globular proteins. Herein, using a range of microscopic and spectroscopic techniques, we show that the LSPT of a functional globular protein, human serum albumin (HSA) is exclusively driven by spontaneous coalescence of liquid-like droplets involving various transient intermediates in a temporal manner. We show that inter-droplet communication via coalescence is essential for both nucleation and growth of amorphous aggregates within individual droplets, which subsequently transform to amyloid-like fibrils. Immobilized droplets neither show any nucleation nor any growth upon aging. Moreover, we found that exchange of materials with the dilute dispersed phase has negligible influence on the LSPT of HSA. Notably, binding of small ligands modulates the feasibility and kinetics of LSPT of HSA, suggesting a possible regulatory mechanism that cells utilize to control the dynamics of LSPT. Further, using a dynamic heterogeneous droplet assembly of two functional proteins, HSA and transferrin (Tf), we show an intriguing phenomenon within the fused droplets where both liquid-like and solid-like phases co-exist within the same droplet, which eventually transform to a mixed fibrillar assembly. These microscopic insights not only highlight the importance of inter-droplet interactions behind the LSPT of biomolecules but also showcase its adverse effect on the structure and function of other functional proteins in a crowded and heterogeneous protein assembly.

## Introduction

Biomolecular condensation of proteins and nucleic acids *via* liquid-liquid phase separation (LLPS) is believed to be responsible for the formation of various membraneless cellular organelles, including stress granules, P-bodies, Cajal bodies, gram granules, nucleolus, centrosomes, and so on.^1,2^ These organelles are actively involved in the spatiotemporal regulation of a variety of critical cellular processes such as signalling, stress regulation, sensing, immune response, noise buffering, genome organization, RNA processing, and so forth.^1,2^ On the other hand, their dysfunctions are associated with various pathological disorders. Therefore, understanding the biomolecular condensation and structure-function relationships of these membraneless compartments are not only important for fundamental research but also crucial for therapeutic reasons.^3^

In this context, tremendous efforts have been made in the last decade to understand the biomolecular condensation of various proteins under physiological conditions.^4–21^ While the presence of intrinsically disorder regions (IDRs) with low complexity domains (LCDs) has been proposed to be a prerequisite for the LLPS of many disease-associated proteins, recent studies have shown that even functional globular proteins lacking any IDRs and/or LCDs can also undergo condensation via LLPS under physiological conditions.^22–27^ These studies revealed that the presence of multivalent non-covalent protein-protein interactions such as electrostatic, hydrophobic, π–π, dipole–dipole, hydrogen bonding, and/or cation–π interactions promotes homotypic LLPS to yield dynamic liquid-like biomolecular condensates/droplets. Our recent efforts have established that the stability and activity of functional enzymes can also be modulated remarkably in a spatiotemporal manner upon biomolecular condensation, indicating its functional role in cellular physiology.^24,27^ In this context, the compartmentalization effect within various synthetic membraneless compartments has been utilized in recent times to boost many chemical and biochemical reactions.^28–31^ Importantly, the heterogeneous and crowded cellular environments in combination with intracellular machineries tightly regulate the complex network of biochemical reactions *via* time-dependent assembly and disassembly of liquid-like biomolecular condensates.^27^

Apart from their functional aspects, recent studies also revealed that liquid-like droplets also act as metastable intermediates during the aberrant liquid-to-solid-like phase transition (LSPT) of many proteins.^8–12,14,19,32–36^ Specifically, LSPT of a wide range of disordered proteins is linked to various neurodegenerative diseases including Alzheimer’s disease, Parkinson’s disease, and amyotrophic lateral sclerosis (ALS). It has been shown that the formation of liquid-like droplets precedes the LSPT during pathological amyloid-like fibrillation via a nucleation and growth-dependent aggregation pathway.^8–12,14,32–36^ The slow nucleation stage is associated with the formation of aggregation prone nuclei which subsequently undergo fast growth/elongation in the presence of monomers and/or oligomers. Moreover, the LSPT via metastable droplets is characterized by gradual change of the viscoelastic nature of the liquid-like droplets along with the concomitant conformational changes of the phase-separated proteins.^37^ These structural changes result in the loss of liquid-like dynamic features into a more rigid gel-like state. However, the origin and nature of the initial microscopic intermediates formed in the nucleation stage during the LSPT of liquid-like droplets remain elusive (Scheme 1). For instant, proteins such as tau,^6^ hnRNPA1,^33^ and FUS^14^ have been shown to undergo LSPT via nucleation at the droplet interface/surface and upon maturation they form a shell of polymerized proteins with a distinct rim and starburst structures (Scheme 1A). Similarly, using a range of chimeric peptides, Winter and coworkers have shown that early fibrils nucleate at the droplet/bulk interface through the formation of single bent fibers.^38^ These studies revealed that the fibril formation does not occur homogeneously inside the droplet phase rather promoted at the interface of the droplets due to the higher surface energy at the interface. On the other hand, initial nucleation and fibril formation for TDP-43^32^ and *α*-Syn^8^ droplets have been shown to originate within the bulk of the droplets rather than at the surface during their LSPT (Scheme 1A). Moreover, it is not clear whether exchange of materials (monomeric or oligomeric proteins) with the surrounding dilute phase has any role in the nucleation and growth of the fibrils during LSPT of liquid-like droplets or the growth is solely regulated by the condensed proteins within the droplet phase (Scheme 1B).

**Scheme 1.**
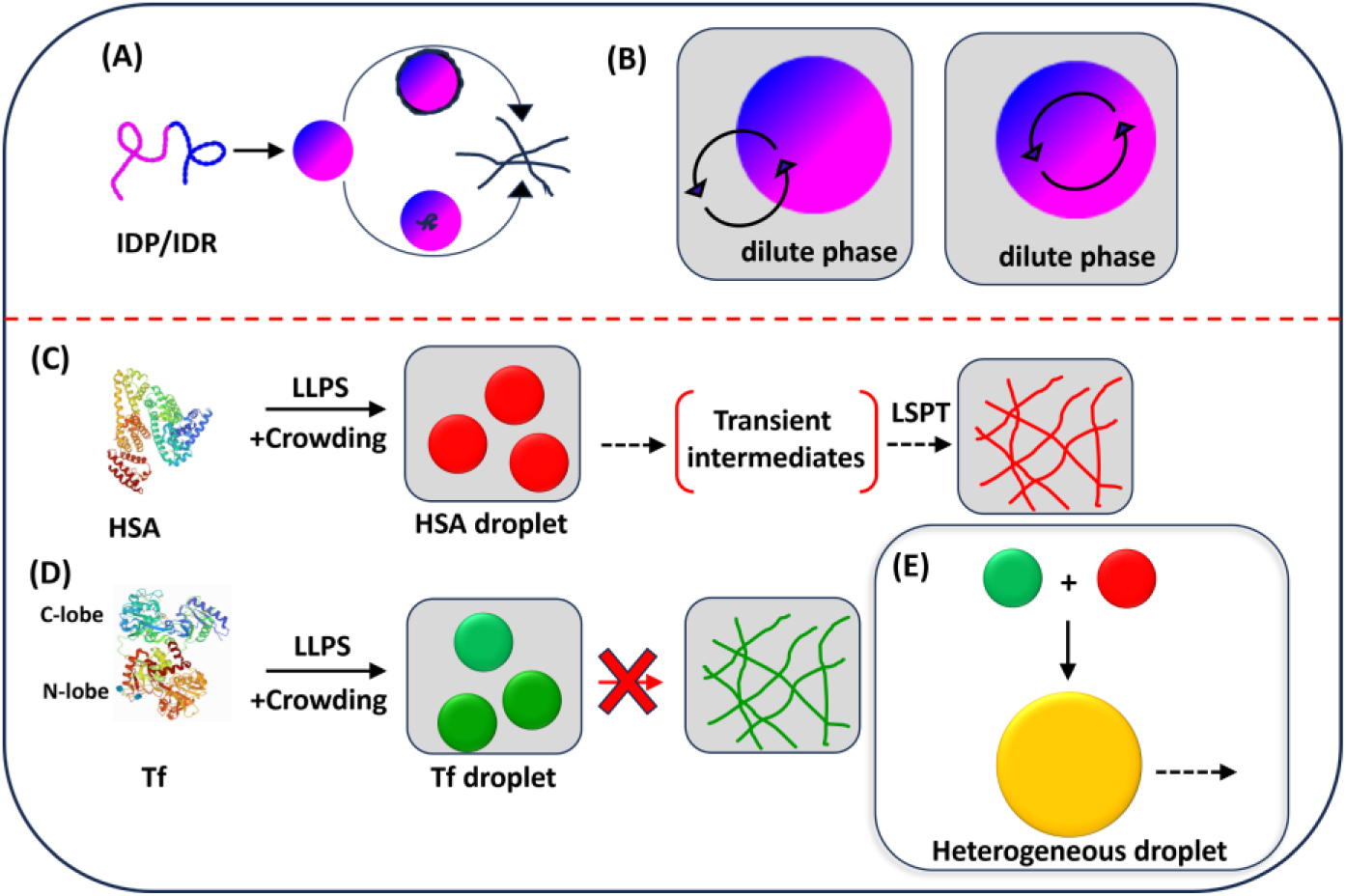
Schematics Showing the (A) Transient Intermediates During Nucleation and LSPT of IDP/IDR Involving Liquid-Like Droplets, (B) Maturation of Droplets With or Without Exchange of Materials with the Dilute Phase, LLPS and LSPT of (C) HSA and (D) Tf, and (E) Coalescence-Driven Formation of Heterogeneous Droplet Assembly from HSA and Tf Droplets.

While much attention was devoted toward understanding the inherent aggregation pathways associated with the LSPT of a wide range of disordered proteins, the microscopic events associated with the LSPT of functional globular proteins during the gradual maturation of the initial transient intermediates remain elusive till date. Understanding these microscopic events and their regulatory factors are particularly important to avoid dysfunction of the associated cellular processes. Notably, the field concerning the LLPS and LSPT of functional globular proteins is still in its infancy stage. In the present study, we aim to address these fundamental questions by studying the LSPT of two functional proteins namely, human serum albumin (HSA) and human serum transferrin (Tf).

Previously, we have demonstrated that both HSA and Tf undergo LLPS upon macromolecular crowding via formation of liquid-like droplets through the involvement of multivalent hydrophobic protein-protein interactions.^23,25^ HSA is the most abundant plasma protein (35–50 g/L) with a molecular weight of ∼66 kDa and involved in transporting various molecules to different parts of the body.^39^ On the other hand, Tf is an iron transport glycoprotein with a molecular weight of 80 kDa, which is synthesized in the liver and subsequently secreted into the plasma.^40^ Uncontrolled aggregation of Tf has been linked with irregular iron distribution and iron overload, which may lead to serious pathological consequences.^40,41^ While the liquid-like droplets of HSA undergo LSPT upon maturation (Scheme 1C),^23^ the liquid-like droplets of Tf do not undergo LSPT (Scheme 1D).^25^ Till date, no systematic efforts have been made to explore the impact of LSPT of a given protein on the physiochemical properties of other nearby functional proteins in a heterogeneous dynamic droplet assembly. We have undertaken the present study to further investigate the early microscopic events associated with the LSPT of HSA in the absence and presence of Tf droplets using a range of microscopic and spectroscopic techniques, including confocal laser scanning microscopy (CLSM) and single droplet Raman spectroscopy. Further, a multicomponent heterogeneous droplet assembly has been designed by utilizing spontaneous coalescence phenomenon of liquid-like droplets of HSA and Tf to understand the phase behavior inside the highly crowded and heterogeneous environment (Scheme 1E).

## Experimental Section

### Materials

Human serum albumin (HSA), human serum transferrin (Tf), polyethylene glycol 8000 (PEG 8000), rhodamine B isothiocyanate (RBITC), fluorescein-5-isothiocyanate (FITC), Hellmanex III, and the Pur-A-Lyzert dialysis kit (molecular weight cutoff 3.5 kDa), thioflavin T (ThT), cetyltrimethylammonium bromide (CTAB), muscovite mica were purchased from Sigma-Aldrich. Sodium dihydrogen phosphate monohydrate (NaH_2_PO_4_.H_2_O), di-sodium hydrogen phosphate heptahydrate (Na_2_HPO_4_.7H_2_O), copper chloride (CuCl_2_), sodium dodecyl sulphate (SDS), sodium azide (NaN_3_), sodium chloride (NaCl), and ethanol (EtOH) were purchased from Merck. 1,2-dipalmitoyl-sn-glycero-3-phosphocholine (DPPC) was purchased from TCI. All the chemicals were used without any further purification. Eco Testr pH1 pH meter was used to adjust the final pH (± 0.1) of all the buffer solutions. Milli-Q water was obtained from a Millipore water purifier system (Milli-Q integral).

### Characterization Techniques

#### Confocal Laser Scanning Microscopy (CLSM)

The confocal images were obtained using an inverted confocal microscope, Olympus fluoView (model FV1200MPE, IX-83) through an oil immersion objective (100×1.4 NA). The samples were excited using two different diode lasers (488 and 559 nm) by using appropriate dichroic and emission filters in the optical path. Confocal images for surface immobilized droplets were captured with drop-cast samples (∼20 μL) onto a cleaned coverslip. The coverslips were dried overnight inside a desiccator before confocal measurement. For liquid-phase experiments, a 20 μL aliquot of the sample solution was drop-cast onto a cleaned glass slide and sandwiched with a Blue Star coverslip. The edges of the coverslips were sealed with a minimal amount of commercially available nail paint.

#### Fourier-Transform Infrared (FTIR) Spectroscopy

FTIR measurements were performed to determine the secondary structure of the HSA using a Bruker spectrometer (Tensor-27). 10 µL of liquid samples of 500 µM HSA in the absence and presence of 10% PEG 8000 in pH 7.4 PBS were used for FTIR measurement. The spectra were recorded in the range of 4000–400 cm^−1^. Fourier self-deconvolution (FSD) method was used to deconvolute the spectra corresponding to the wavenumbers 1700−1600 cm^−1^.^25^ The Lorentzian curve fitting was done to fit the spectra using origin 8.1 software. The experiments were performed twice with similar observations.

#### UV−vis Spectrophotometry

Absorption spectra were recorded in a quartz cuvette (1 cm × 1 cm) using a Varian Carry 100 Bio UV−visible spectrophotometer. *Fluorescence Spectroscopy.* The fluorescence spectra were recorded in a quartz cuvette (1 cm × 1 cm) using a HORIBA Jobin Yvon, model FM-100 Fluoromax-4 Spectrofluorometer at constant temperature 37 °C. The slit width was kept at 3 nm.

#### Circular Dichroism (CD) Spectroscopy

CD spectra were recorded on a JASCO J-815 CD spectropolarimeter using a quartz cell of 1 mm path length with a scan range of 190-260 nm. Scans were recorded with a slit width of 1 mm and speed of 50 nm/min. For CD measurements, protein solutions were diluted by 100-fold to make the final working concentration of 5 μM. The mean residue ellipticity (MRE) in deg cm^2^ dmol^-1^ of HSA at 222 nm was calculated using the formula,

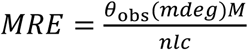

where *Ɵ*_obs_ is the CD in millidegrees, *M* is the molecular weight of the protein in g dmol^-1^, *n* is the number of amino acid residues (585 for HSA), *l* is the path length (0.1 cm) of the cuvette and *c* is the concentration of the protein in gL^-1^.

#### Laser Raman Spectroscopy

Raman spectra of protein samples were recorded using a Horiba–Jobin Yvon micro-Raman spectrometer equipped with a 532 nm excitation laser and a CCD detector. For Raman measurements of protein droplets, the PEG molecules were removed by centrifugation. The resuspended dense phase was deposited onto a glass slide covered with an aluminium sheet. A 532 nm excitation laser with 40 mW (100%) laser power was used for excitation with an exposure time of 10 s. The laser was focused using a 100× objective lens (Nikon, Japan). The spot size of the laser beam in the Raman spectrometer was ∼1 μm, and the diffraction grating of 600 lines/mm was used to disperse the scattering light. Data were collected in the backscattering mode with flat correction using an inbuilt method in Raman spectrometer software. The obtained experimental spectra were baseline corrected and normalized with respect to the phenylalanine ring breathing band at 1005 cm^−1^ using origin 8.1 software. All the recorded spectra were averaged over 3 to 5 scans.

#### Atomic Force Microscopy

AFM images were acquired using a PERK atomic force microscope operating in tapping mode. For sample preparation, 10 μL aliquots were taken and deposited onto freshly cleaved muscovite mica. XEI software was used for image processing. The height profiles were analysed from XEI.

#### Rheology Measurement

Viscoelastic characterization of the HSA gel was carried out by rheology. Anton Paar Physica rheometer (model no MCR 301) was used for all rheological experiments. HSA gel was placed on the rheometer disc by using a micropipette. A parallel plate with a diameter of 25 mm was used for this experiment. The TruGap was set at 0.5 mm. The mechanical strengths of the HSA gel were determined by strain sweep experiment at a constant frequency (10 rad s^−1^). In the strain sweep experiment, the storage modulus (G’) and loss modulus (G’’) were plotted as a function of strain (%).

## Results and Discussion

### LSPT and Material Properties

We first investigated the feasibility of LSPT and associated changes in the material properties of aqueous solution of HSA and Tf in the presence of 10% PEG as a macromolecular crowder under physiological conditions. The aqueous solution of 500 µM HSA in the presence of 10% PEG in pH 7.4 PBS was equilibrated at 37 ℃ for a period of 1–14 days. The daylight photographs of aqueous solution of HSA revealed time-dependent transition from liquid to solid-like gel state upon 14 days of aging at 37 ℃ (Figure 1A). The turbidity measurement revealed gradual increase in the turbidity up to 7 days of aging (Figure S1), beyond which the aqueous solution gradually transformed into a semi-solid gel-like state. In contrast, the aqueous solution of 1 μM Tf in the presence of 10% PEG remains isotropic in nature over a period of 30 days without any LSPT (Figure 1B). These findings are consistent with our previous reports.^23,25^ The rich phase behavior of HSA suggests time-dependent changes of its material properties. To gain further understanding of the inherent material properties of solid-like gel state of HSA, we performed rheological measurements. The strain-sweep study of the solid-like gel states at d-11 and d-14 showed that the values of the elastic storage modulus G’ were significantly higher than the loss modulus G” up to a strain of 125% at d-11, and 551% at d-14, suggesting gradual increase in the solid-like elastic behavior over the liquid-like viscous properties upon aging (Figure 1C).^42^ These changes in the material properties were accompanied by progressive alterations in the secondary structure of HSA as revealed from the circular dichroism study (Figures 1D and S2).

**Figure 1.**
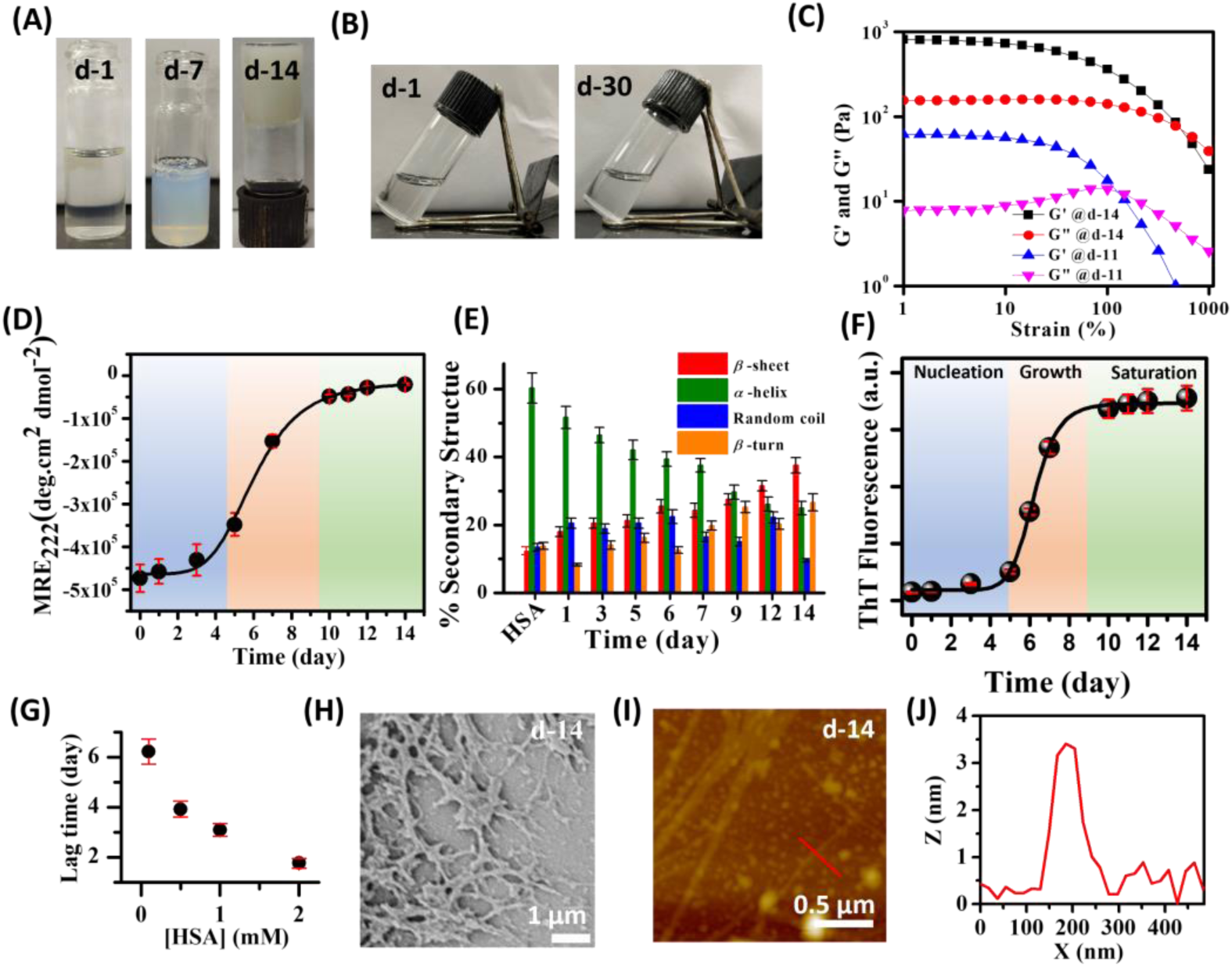
Time-dependent daylight photographs of aqueous solution of (A) 500 µM HSA and (B) 1 µM Tf in the presence of 10% PEG. (C) Rheological strain-sweep study of 500 µM HSA in the presence of 10% PEG at d-11 and d-14. (D) Changes in the MRE values at 222 nm of 500 μM HSA in the presence of 10% PEG (diluted by 100-fold) over a period of 14 days. (E) Changes in the secondary structures of HSA in the presence of 10% PEG estimated from the deconvoluted FTIR spectra upon aging a period of 14 days. (F) ThT-fluorescence assay in the presence of 500 μM HSA and 10% PEG over a period of 14 days. (G) Changes in the lag time as a function of HSA concentrations. (H) FESEM and (I) AFM images showing fibrillar assemblies of HSA at d-14. (J) AFM height profile of HSA fibril at d-14. Data represent mean ± s.e.m. for three independent measurements (*n* = 3).

The time-dependent changes in the mean residual ellipticity (MRE) values at 222 nm revealed a typical sigmoidal feature with a sharp transition beyond 5 days of aging, which saturates beyond 12 days, suggesting a gradual decrease in the *α*-helix content of HSA.^43,44^ To further validate this observation, we recorded the FTIR spectra as a function of aging and quantitatively estimated the secondary structure content of HSA (Figures 1E and S3). The recorded FTIR spectra were deconvoluted and the secondary structures were estimated using the area under the deconvoluted peaks (Figure S4). The estimated secondary structure of HSA revealed 61.0% *α*-helix, 12.1% *β*-sheet, 13.1% random coil, and 13.8% *β*-turn in the absence of any crowders. A continuous decrease in the *α*-helix content with a concomitant increase in the *β*-sheet and *β*-turn contents was noticed. At d-14, a significantly altered conformation with 25% *α*-helix, 38% *β*-sheet, 9.9% random coil, and 27.1% *β*-turn was observed (Figure 1E). These data suggest that the LSPT of HSA is characterized by a decrease in the *α*-helix content and concomitant increase in the *β*-sheet and *β*-turn contents. Similar LSPT via formation of *β*-sheet rich amyloid-like fibrils has been reported previously for many disordered proteins.^4,6,8–12^ To know whether these aggregates of HSA are linked to amyloid-like *β*-sheet structures, we performed thioflavin T (ThT) assay. ThT is a well-known fluorescent marker for amyloid-like structures and exhibits enhanced fluorescence upon binding with the *β*-sheet structures.^45,46^ The aggregation kinetics of HSA were monitored by measuring the fluorescence intensity of 20 μM ThT in the presence of 500 μM HSA and 10% PEG. A classical sigmoidal curve with an initial lag phase (nucleation), growth phase (elongation), and saturation phase was observed upon 14 days of aging (Figure 1F). Importantly, the lag time was found to be dependent on the initial concentration of HSA used (Figures 1G and S5), suggesting a nucleation-dependent polymerization mechanism similar to those observed previously for several other proteins.^32,47–49^ Importantly, a 22-fold enhancement in the fluorescence intensity of ThT was observed at d-14 relative to d-1 (Figure S6), suggesting gradual maturation of initial native protein into a more compact amyloid-like aggregates. FESEM and AFM measurements revealed the presence of distinct fibrils having diameter of ∼3.4 nm with several microns in length (Figure 1H–J). These findings reveal that while HSA undergo spontaneous LSPT via formation of amyloid-like fibrils, Tf failed to undergo LSPT even after a prolong time of aging under physiological conditions. To gain further insight into the microscopic events associated with the observed LSPT of HSA, we directly visualized the transient intermediates under the confocal microscope.

### Transient Intermediates During LSPT of HSA

To visualize various transient intermediates during the LSPT of HSA, we performed CLSM imaging using rhodamine B isothiocyanate (RBITC)-labeled HSA. For confocal imaging, an aqueous solution of 500 μM RBITC-labeled HSA in the presence of 10% PEG was incubated at 37 ℃ in 50 mM pH 7.4 PBS, and at different time intervals (1–14 days), 20 μL aliquot of the sample solution was placed inside a sealed liquid chamber (Figure 2A). Confocal fluorescence image of the d-1 sample revealed the presence of well-dispersed droplets (Figure 2B), which exhibit characteristic liquid-like properties such as fusion/coalescence, dripping, and surface wetting (Figure S7). These results are consistent with our previous report and suggest the formation of liquid-like droplets via homotypic LLPS of HSA via multivalent hydrophobic protein-protein interactions.^23^ It should be noted that upon LLPS, HSA formed liquid-like droplets instantaneously within few minutes upon macromolecular crowding (Figure S8). While these droplets remained spherical in shape upon aging up to d-5 (Figure 2B), the mean size of the droplets increased from 3.59 ± 0.18 µm at d-1 to 6.17 ± 0.34 µm at d-5 (Figure 2C). A closer look at the d-5 confocal image revealed the presence of several bright localized fluorescence spots within the droplets (Figure 2B, yellow arrows). Over time, we observed increase in the number of these bright fluorescence spots as well as their size within the droplet phase between d-5 and d-6 (Figure S9).

**Figure 2.**
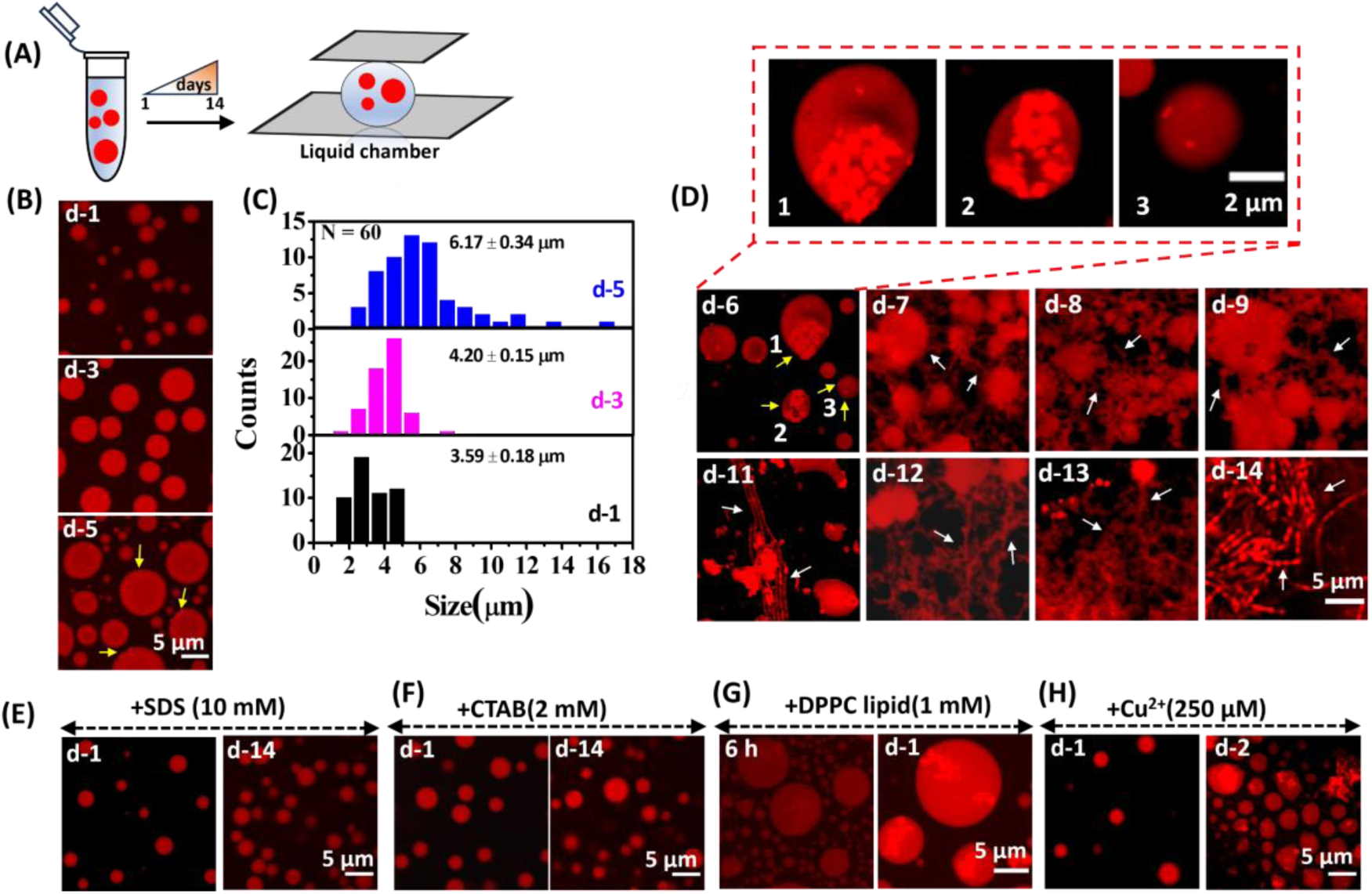
(A) Schematic showing the liquid chamber configuration for confocal imaging. (B) CLSM images and (C) size distribution histograms of RBITC-labeled HSA droplets as a function of aging over a period of 5 days. Data represent mean ± s.e.m for three independent measurements (*n* = 3). (D) CLSM images of RBITC-labeled HSA droplets from d-6 to d-14. The enlarged CLSM images show the presence of initial nuclei and amorphous aggregates within the representative droplets at d-6. CLSM images of RBITC-labeled HSA droplets in the presence of (E) 10 mM SDS, (F) 2 mM CTAB, (G) 1 mM DPPC lipid, and (H) 250 µM Ca^2+^ at different time intervals.

Moreover, these initial clusters were mainly originated at the edges/interfaces of the spherical droplets and undergo random diffusion within the droplet boundaries. Notably, the appearance of these clusters at d-5 exactly coincides with the onset of the sharp transition observed for MRE value and ThT emission. Therefore, we assigned these initial clusters as aggregation competent nuclei of HSA, which originated at the droplet interfaces possibly due to the high surface energy at the interface.^6,14,33,38^ At d-6, we observed the presence of initial nuclei as well as higher order aggregates within the individual droplets having bright localized fluorescence signals (Figure 2D, yellow arrows). While higher order amorphous aggregates were mainly observed inside the larger sized droplets, smaller sized droplets mainly revealed the presence of short-nucleated structures. Upon further aging for a period of d-7 to d-9, the amorphous aggregates of HSA spread beyond the boundaries of individual droplets and transformed into inter-droplet networks (Figure 2D, white arrows). Notably, the mobility of these matured aggregates reduced significantly at d-9 compared to that at d-6, signifying the on-set of LSPT. The number of spherical droplets decreased significantly beyond d-11 and long fibrillar networks emerged. The CLSM image at d-14 revealed a complete conversion of the spherical droplets into long matured fibrils (Figure 2D), which is consistent with our FESEM and AFM results. Hence, these microscopic observations authenticate that the liquid-like droplets of HSA act as dynamic metastable scaffolds to host various transient dynamic intermediates during the slow and spontaneous LSPT via nucleation and growth-mediated aggregation pathway. To know whether exchange of materials (monomers and/or oligomers) with the dilute dispersed phase has any role in the observed aggregation pathway, we performed phase separation assays after replacing the dilute dispersed phase with equal volume of PBS buffer containing 10% PEG. Notably, we observed similar time-dependent nucleation and growth of HSA fibrils even in the absence of dilute protein phase (Figure S10), indicating that only the phase-separated proteins inside the droplet phase take part in the aggregation process and the exchange of materials with the dilute dispersed phase has negligible impact on the aggregation process.

### Ligand Binding Regulates the LSPT of HSA

Ligands are known to influence the feasibility of LLPS of scaffold proteins.^50^ To know the effect of ligand binding on the LLPS and LSPT of HSA, we utilized a variety of potential ligands, namely surfactants, lipids, and metal ions and performed the LSPT assays using CLSM under similar experimental conditions. These additives are known to bind with serum albumins at physiological conditions.^51–54^ Interestingly, the presence of 10 mM negatively charged sodium dodecyl sulfate (SDS), and 2 mM positively charged cetyltrimethylammonium bromide (CTAB) had negligible impact on the LLPS of HSA; however, both these surfactants inhibited the LSPT of HSA droplets. Neither any nucleation nor any fibrillar growth of HSA was observed in the presence of both SDS and CTAB over a period of 14 days (Figure 2E,F). In addition, the mean droplet size did not change significantly even after 14 days of aging in the presence of SDS and CTAB (Figure S11), suggesting inhibition of the spontaneous coalescence of droplets in the presence of these surfactants. Interestingly, we noticed that while many droplets collided with neighbouring droplets, majority of them failed to undergo coalescence in the presence of 10 mM SDS at d-5. Earlier, by using coarse-grained simulations, Espinosa and coworkers demonstrated size conservation in biomolecular condensates in the presence of surfactant clients by considering droplet surface tension.^55^ Recently, Arosio and coworkers observed similar inhibition of LSPT of hnRNPA1 droplets in the presence of 0.03% SDS.^33^ Using a *β*-peptide (1–28), Zagorski and coworkers showed that membrane-like charged surfaces are effective in stabilizing the soluble *α*-helical conformation of the *β*-peptide in the physiological pH range and prevent aggregation into the *β*-sheet.^56^ It is important to mention that at lower concentration of surfactant, we indeed observed delayed nucleation within the droplets (Figure S12), suggesting that the inhibition effect is concentration dependent. In contrary, the presence of either 1.0 mM zwitterionic DPPC, 250 μM Cu^2+^, or 10 mM Ca^2+^ was found to promote the early nucleation and aggregation of HSA (Figures 2G,H and S13). Further, a remarkable increase in the mean droplet size was noticed in the presence of DPPC and Cu^2+^ (Figure S14). These findings indicate that ligand binding effectively regulates the intermolecular protein-protein interactions as well as inter-droplet interactions required for nucleation and growth of HSA fibrils. We hypothesized that this regulation might be due to the conformational alteration of the ligand-bound HSA which modulates the favourable protein-protein contacts. To validate this argument, we monitored the secondary structure of HSA in the absence and presence of various ligands using CD and FTIR measurements (Figure S15,16) A noticeable decrease in the *α*-helix content with concomitant increase in the *β*-sheet content was observed in the presence of all the ligands, suggesting binding of these ligands with the protein residues. This indicates that the unusual stabilization against aberrant aggregation of HSA observed in the presence of charged surfactants must be due to some additional factors that stabilize the phase-separated droplets. Notably, charged surfactants are known to bind at the aqueous interfaces and lower the interfacial surface tension.^33,55^ We propose that the additional layer of charged surfactants at the surface of these droplets provides an electrostatic barrier against spontaneous coalescence and at the same time lowers the surface tension of the droplets, which effectively prevent the aberrant protein aggregation. Therefore, our data suggest that the presence of these ligands not only alters the secondary structures of phase-separated proteins, but also modifies the surface properties of liquid-like droplets. We believe that living cells might also utilize similar regulatory machineries to control the feasibility of LSPT of biomolecules to avoid dysfunction of critical cellular processes.

### Surface Immobilization of HSA Droplets

To know whether immobilized droplets able to undergo LSPT via similar transient intermediates upon maturation, we immobilized the liquid-like droplets of HSA on cleaned glass coverslips and aged for a period of 1–14 days (Figure 3A). CLSM images revealed no noticeable change in the droplet morphology upon immobilization even after 14 days of aging (Figure 3B).

**Figure 3.**
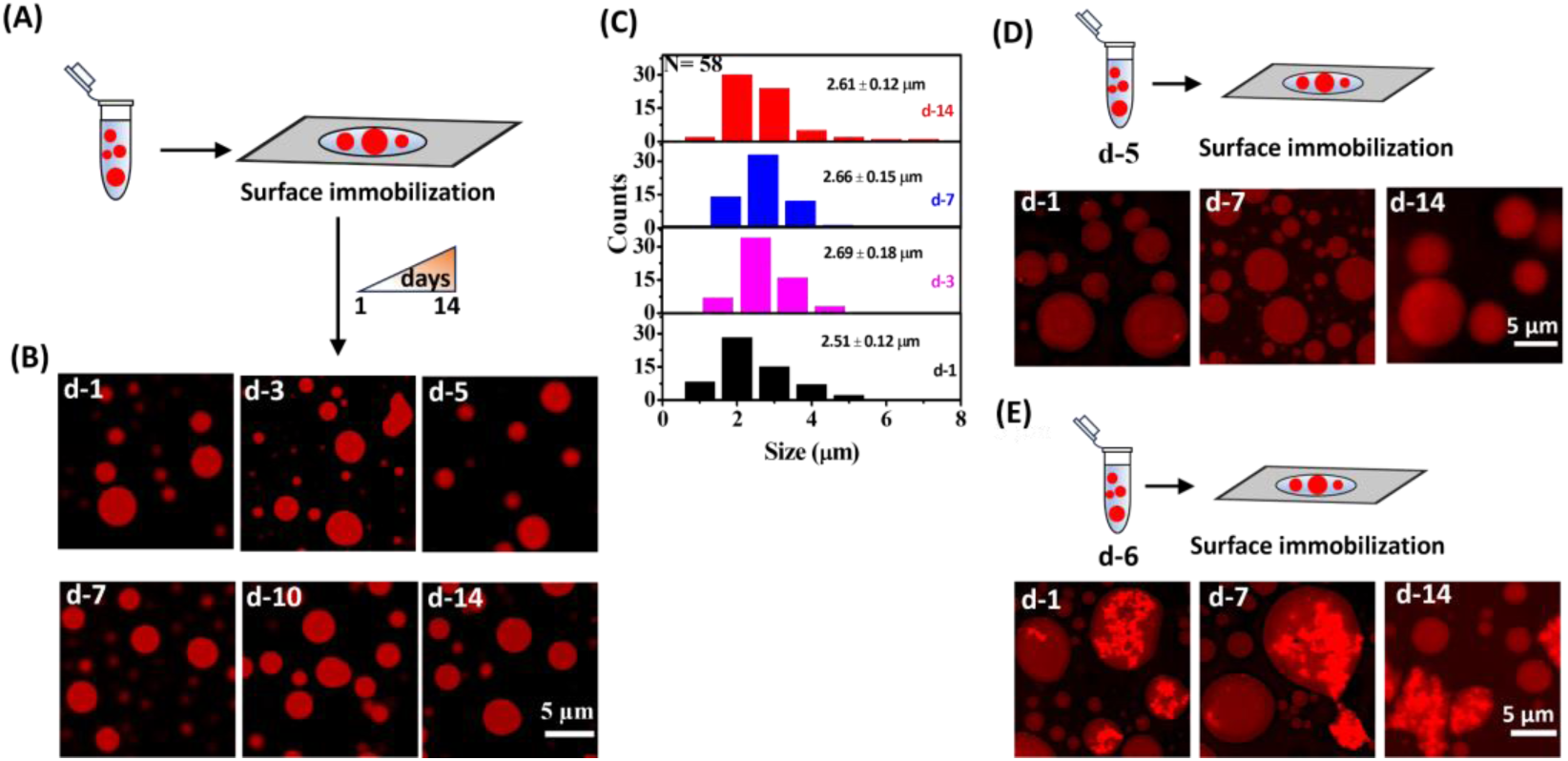
(A) Schematic showing the surface immobilization of HSA droplets on the glass surface. (B) CLSM images of RBITC-labeled HSA droplets as a function of surface aging for a period of 14 days. (C) Size distribution histograms of the surface-aged HSA droplets at different time intervals. Data represent mean ± s.e.m. for three independent measurements (*n* = 3). Confocal images of (D) d-5 and (E) d-6 solution aged HSA droplets at different time intervals after surface immobilization.

Further, the mean size of these immobilized droplets remained almost unaltered upon surface aging (Figure 3C), suggesting lack of any coalescence events. These observations indicate that in the absence of any coalescence, immobilized droplets failed to undergo LSPT. Therefore, our study reveals that mere formation of liquid-like droplet is not enough to trigger aberrant aggregation and LSPT in the absence of coalescence. To check whether droplets with pre-formed nuclei able to mature over time, we first equilibrated HSA droplets in the solution phase for 5 and 6 days and then immobilized them on the glass coverslips (Figure 3D,E). However, both d-5 and d-6 samples failed to mature into fibrillar network over time irrespective of the extent of nucleation in individual droplets. This suggests that the growth of the initially formed nucleated structures inside the immobilized droplets inhibited completely in the absence of coalescence. Therefore, our findings authenticate the essential role of coalescence behind the nucleation and growth of aggregates inside the phase-separated droplets of HSA during the LSPT.

### Coalescence-Driven Enrichment of Proteins Inside Individual Droplets

Upon establishing the essential role of coalescence behind the nucleation and aggregation of HSA, we wanted to know whether increased protein concentration within the individual droplets is responsible for the coalescence-driven enhanced protein-protein interactions. Initially, we quantitatively estimated the protein concentration in the dense droplet phase (*C*_D_) and dilute supernatant phase (*C*_S_) of the d-1 aged sample using the conventional centrifugation method.^10^ The dilute supernatant phase was carefully separated from the dense droplet phase after centrifugation at ∼18,000 rpm for 30 min. After appropriate dilution with denaturing buffer, the concentration of HSA in the dense droplet phase and dilute phase was estimated from the UV-vis spectrophotometry using a molar extinction coefficient of 35,700 M^-1^cm^-1^ at 280 nm.^57^ The concentration of HSA in the droplet and dilute dispersed phase was found to be 15.6 and 0.19 mM, respectively (Figure 4A). This 82-fold enrichment in the protein concentration inside the droplet phase is consistent with previous reports on various other phase-separated proteins.^6,9,10,33^

**Figure 4.**
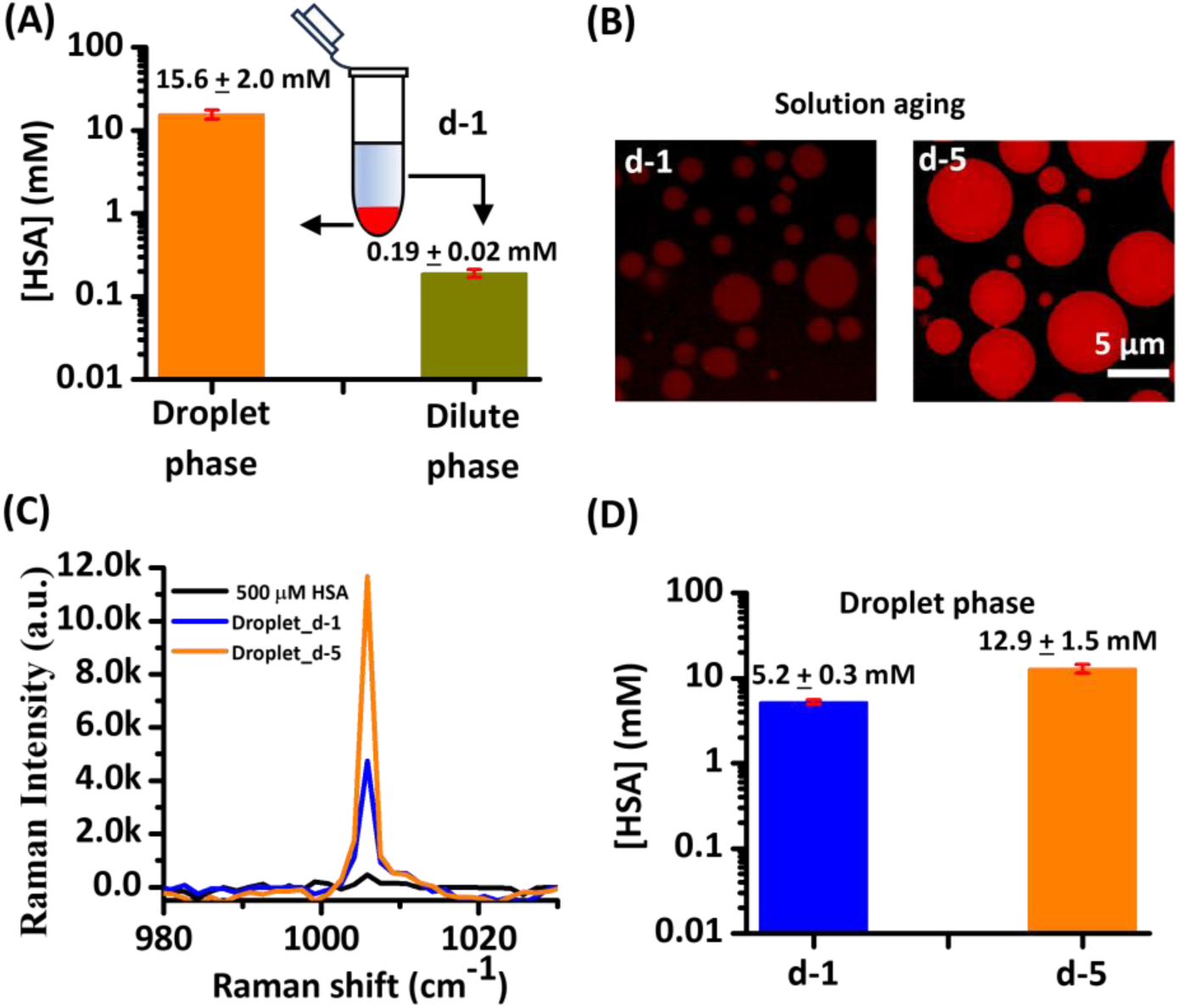
(A) Bulk protein concentrations in droplet and dilute phases estimated using UV-vis spectrophotometry. Data represent mean ± s.e.m. for three independent experiments (*n* = 3). (B) CLSM images of the solution-aged HSA droplets at d-1 and d-5. (C) Single droplet vibrational Raman spectra of 500 µM HSA and HSA droplets at d-1 and d-5 averaged over five independent runs (*n* = 5 samples). (D) Estimated protein concentrations within the droplets (n = 5) at d-1 and d-5 obtained from the relative Raman intensity at 1005 cm^-1^. Data represent mean ± s.e.m (*n* = 5 droplets).

Next, to know the effect of aging on the local protein concentration within the individual droplets, we visualized the changes in the relative fluorescence intensity of RBITC-labeled HSA droplets at d-1 and d-5 under the same instrumental and experimental settings. CLSM images revealed significant increase in the size as well as relative fluorescence intensity within the d-5 aged droplets compared to that in d-1 aged droplets (Figure 4B). In contrast, no such increase in the size as well as in the fluorescence signal was noticed for surface immobilized droplets even after 7 days of surface aging under similar conditions (Figure S17). These data thus suggest coalescence-driven increase in the local protein concentration within individual droplets upon solution phase aging.

We quantitatively estimated the local protein concentration within the individual droplets as a function of aging using a label-free Raman spectroscopy method *via* monitoring the characteristic phenylalanine Raman peak intensity at 1005 cm^-1^ as a marker.^33^ Figure 4C shows the phenylalanine Raman band of 500 μM dispersed HSA and phase-separated HSA droplets averaged over five different droplets (*n* =5). Notably, a 10.4-fold increase in the Raman intensity at 1005 cm^-1^ was observed upon phase separation at d-1 relative to that of 500 μM dispersed HSA sample. Upon further aging, the Raman intensity increased remarkably by a factor of 25.7-fold at d-5. We estimated the effective protein concentrations within the droplet phase by comparing the Raman intensity of the 500 µM dispersed HSA sample with those of the aged samples. The local protein concentration was found to be 5.2 ± 0.3 and 12.9 ± 1.5 mM for d-1 and d-5 aged samples, respectively (Figure 4D). Clearly, these data authenticate coalescence-driven enrichment of local protein concentration within the individual droplets during solution phase aging. We believe that this enrichment is essential to cross a threshold protein concentration (*C*_t_) required for nucleation and subsequent aggregation within the droplets.

### Single Droplet Raman Spectroscopy

To gain residue level structural information within the individual droplets during the LSPT of HSA, we performed single droplet vibrational Raman spectroscopy with a laser Raman setup equipped with a microscope to visualize individual droplets and fibrils. Vibrational Raman spectroscopy is a very useful and highly sensitive label-free technique as the signal originates exclusively from the local molecular vibrations of the protein residues and therefore, this technique has been utilized extensively in recent times to get the molecular information and conformational heterogeneity of biomolecules.^10,11,58–61^ Raman spectra were recorded by using a micro-Raman spectrometer by focusing a 532 nm excitation laser light source (40 mW) into the individual droplets/fibrils using a 100× objective lens (Figure 5A). The recorded Raman spectra were averaged over three independent runs (n = 3) and the normalized average spectra were plotted as a function of aging for a period of 1– 14 days (Figure 5B).

**Figure 5.**
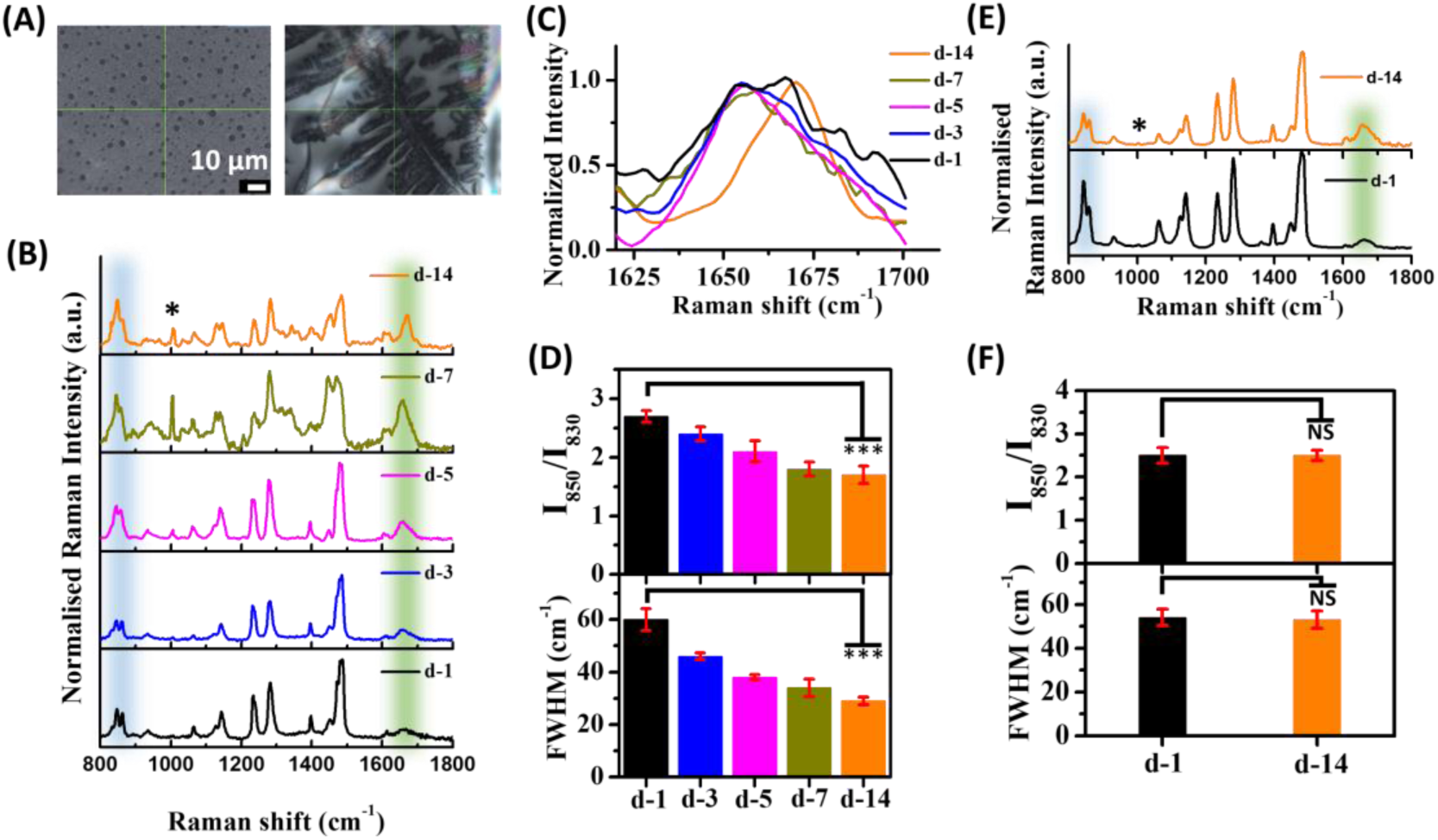
(A) Raman microscopic images showing the focused laser beams (*λ* = 532 nm) on individual droplets and fibrils of HSA. (B) Changes in the single droplet Raman spectra (*λ* = 532 nm) of HSA droplets as a function of aging. (C) Normalized Raman spectra of HSA samples in the amide-I band region as a function of aging. (D) Changes in the FWHM of amide I band and tyrosine Fermi doublet (*I*_850_/*I*_830_) ratio of HSA samples as a function of aging. Data represent mean ± s.e.m. for three independent experiments (*n* = 3). (E) Changes in the single droplet Raman spectra of surface-aged HSA samples. (F) Changes in the FWHM and tyrosine Fermi doublet (*I*_850_/*I*_830_) HSA samples. Data represent mean ± s.e.m. for three independent experiments (*n* = 3). Each single droplet Raman spectrum was averaged over 3 runs (*n* = 3 droplets). Raman spectra were normalized with respect to the phenylalanine peak at 1005 cm^-1^, denoted by an asterisk. Statistical significance was assessed by a two-tailed, unpaired Student’s *t*-test with ***, *P* < 0.001 and not significant (NS), *P* > 0.05.

The Raman spectrum of HSA droplets at d-1 revealed the presence of all the characteristic vibrational modes of the polypeptide chain, including amide I (1630–1700 cm^-1^), amide III (1230–1300 cm^-1^), aromatic residues (tryptophan, tyrosine, and phenylalanine), and other aliphatic side-chain vibrations (Figures 5B, S18 and Table S1). Here, we mainly focused our attention on the spectral features associated with the amide I band and tyrosine Fermi doublet (850 and 830 cm^-1^). The temporal evolution of the amide I band in the region of 1630–1700 cm^-1^ revealed a noticeable change in the peak position and full width at half-maximum (FWHM) upon aging for 14 days. The amide-I band for the d-14 aged sample showed a 14.1 cm^-1^ shift in the higher frequency region relative to those at d-1 and d-7 aged samples (Figure 5C). This increase in the carbonyl (–C=O) stretching frequency of the amide I band suggests a more compact and rigid conformational state of HSA upon maturation.

Further, a gradual decrease in the FWHM of the amide-1 band from a value of 60.0 ± 4.2 cm^-1^ at d-1 to a value of 29.0 ± 1.5 cm^-1^ at d-14 was observed (Figure 5D, lower panel), suggesting a progressive change in the protein conformation from a flexible and heterogeneous conformational state to a more homogeneous and compact conformational state.^10,11,58^ On the other hand, the intensity ratio of the tyrosine Fermi doublet (*I*_850_/*I*_830_) in the Raman spectrum indicates the hydrogen bonding tendency of the surrounding water molecules with the phenolic hydroxyl group of tyrosine and exhibits a value of ≥2.0 for a well hydrated residue. Notably, the ratio of *I*_850_/*I*_830_ decreased from a value of 2.7 ± 0.1 at d-1 to a value of 1.7 ± 0.2 at d-14 (Figure 5D, upper panel), indicating lower extend of hydration upon formation of amyloid-like mature fibrillar network upon aging.^10,11,58^ In contrast, the Raman spectra of immobilized droplets revealed neither any change in the FWHM of amide-I band nor any change in the value of I_850_/I_830_ upon aging (Figure 5E,F), suggesting conformational stability of the immobilized droplets.

### LSPT in Dynamic Heterogeneous Droplet Assembly

One of the primary functions of membraneless cellular organelles is to concentrate various biomolecules inside their confined space. How this crowded and heterogeneous environment affects the behavior of individual biomolecules is a question of our present interest. To mimic this cellular heterogeneity, we designed a dynamic heterogeneous droplet assembly by combining the aggregation prone HSA droplets with structurally robust Tf droplets to explore the impact of LSPT of HSA on the physicochemical properties of Tf. The droplet assembly was prepared by mixing the phase-separated droplets of 500 µM RBITC-labeled HSA and 50 µM FITC-labeled Tf in the presence of 10% PEG in a 1:1 ratio and equilibrated at 37 ℃ for a period of 1–14 days (Figure 6A). Temporal evolution of this heterogeneous assembly was monitored at different time intervals using CLSM. Instantaneously after mixing (within 5 min), we observed distinct green and red emissive spherical droplets of Tf and HSA, respectively (Figure 6B). Importantly, the merged image revealed yellow signals from the fused droplets along with individual red and green emissive droplets. Notably, within 5 days, we observed only fused droplets with distinct yellow signals due to the coalescence events. The merged image at d-5 revealed the presence of aggregated clusters of RBITC-labeled HSA within the fused droplets. This effect was more pronounced in d-6, where distinct aggregates of RBITC-labeled HSA emerged within the fused droplets as revealed from the appearance of bright red emissive aggregated clusters.

**Figure 6.**
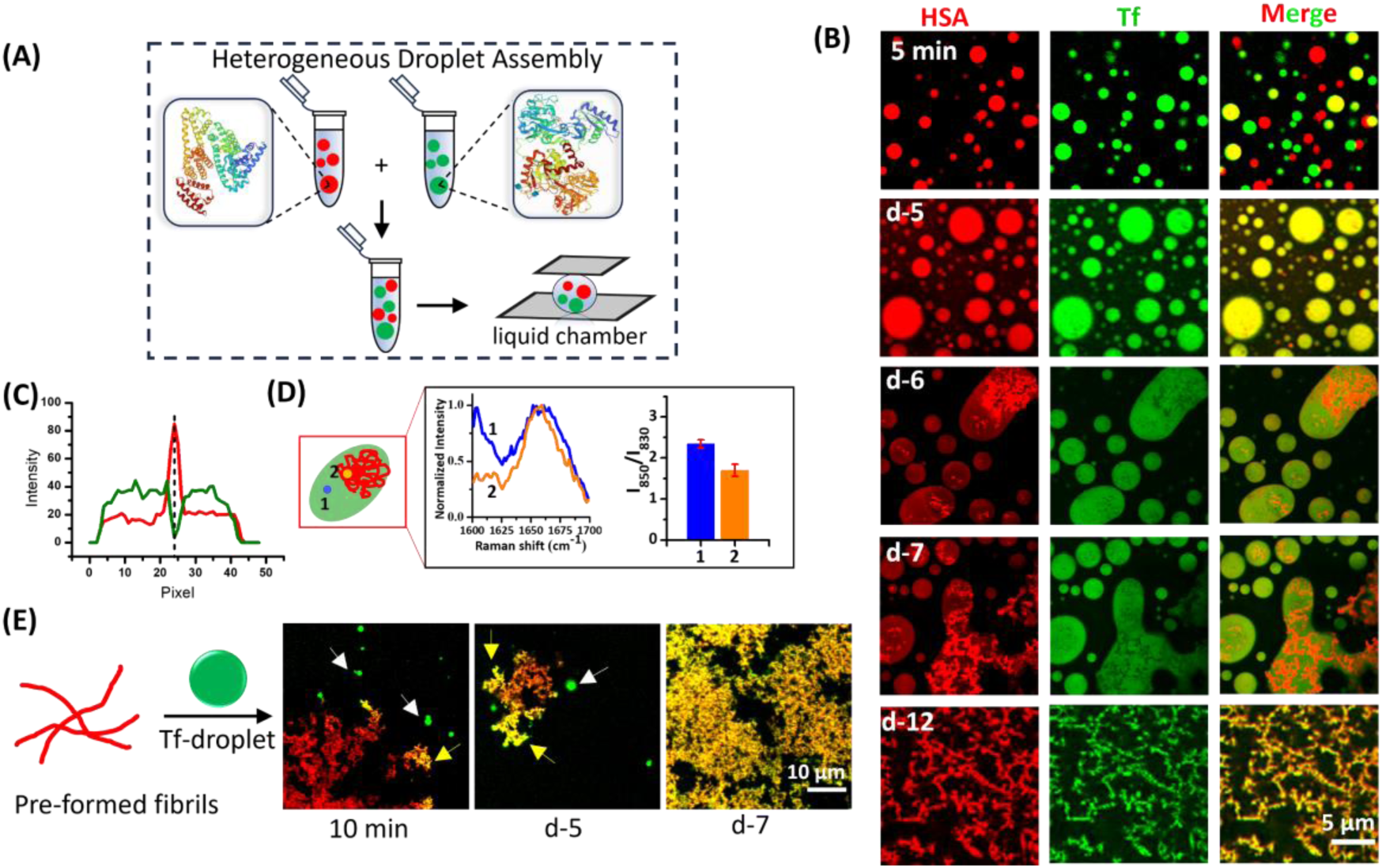
(A) Schematics showing the sample configuration for heterogeneous droplet assembly. (B) CLSM images of heterogeneous droplet assembly at different time intervals. (B) Intensity line plots of green and red channels within a representative fused droplets at d-6. (C) Spatially resolved single droplet Raman features of amide I band and tyrosine Fermi doublet (*I*_850_/*I*_830_) ratio from a heterogeneous fused droplet at d-6. Data represent mean ± s.e.m. for three independent measurements (*n* = 3). (E) CLSM images showing time-dependent adsorption of FITC-labelled Tf droplets (white arrows) on the surface of pre-formed RBITC-labelled HSA fibrils to yield mixed fibrillar assembly (yellow arrows).

Interestingly, the CLSM image in the green channel revealed no such clustering of the green signals, indicating that FITC-labeled Tf do not undergo aggregation within the fused droplets (Figure 6B). However, we observed non-uniform emission from FITC-labeled Tf in the green channel with several dark spots/regions. Notably, the locations of these dark spots/regions in the green channel exactly matched with the locations of intense red emission in the red channel as revealed from the intensity line profiles (Figure 6C). These findings indicate that the dark spots/regions in the green channel originated due to the exclusion of the FITC-labelled Tf by the nucleated/aggregated clusters of RBITC-labelled HSA. At d-7, more extensive aggregation of HSA was observed within the fused droplets without any evidence of aggregation/clustering of Tf. These microscopic findings clearly indicate a contrasting behavior of HSA and Tf within the phase-separated fused droplets. Although, both HSA and Tf undergo LLPS via multivalent hydrophobic interactions,^23,25^ we did not observe any cross-coupled aggregates inside the fused droplets. It is obvious from our findings that the specific homotypic protein-protein interactions dictate the spatial organization of HSA within the heterogeneous droplets. Our data thus demonstrate the dominant role of specific homotypic protein-protein interactions over non-specific heterotypic protein-protein interactions within the fused droplets of HSA and Tf. This intriguing finding is highly relevant in the context of intracellular organelles which are known to concentrate a diverse range of biomolecules and regulate their complex physiological functions inside the heterogeneous environment *via* specific biomolecular interactions.^62^

To selectively interrogate the local environments around the polypeptide chains of HSA and Tf within these heterogeneous fused droplets, we performed vibrational Raman spectroscopy by selectively focusing the laser at different locations within the fused droplets (Figure 6D). The Raman spectrum recorded from a homogeneous part of the droplet revealed an amide I band with a FWHM of 60.0 ± 3.9 cm^-1^. However, the Raman signal collected from a region having aggregated structure within the fused droplets showed an amide I band with a FWHM of 39.0 ± 2.5 cm^-1^. Similarly, the *I*_850_/*I*_830_ ratio from the homogeneous and aggregated structures was found to be 2.3 ± 0.1 and 1.7 ± 0.2, respectively. These spatially resolved Raman features of the fused droplets authenticate that while the polypeptide chain of HSA displays a more rigid and compact conformational state, the well-hydrated polypeptide chain of Tf exhibits a substantial amount of conformational heterogeneity inside the droplets. These contrasting and unique microscopic events within the small volume of fused droplets highlight an important aspect of specific protein-protein interactions and mimic the intracellular organization and functioning of biomolecules inside the membraneless organelles. Considering the half-life of cellular Tf ∼7–8 days,^40,63^ our findings indicate that the amorphous aggregates of HSA do not perturb the native-like conformation of phase-separated Tf within its half-life period. Upon further maturation, distinct amyloid-like fibrillar networks in both red and green channels were observed and resulted in a mixed fibrillar assembly at d-12 (Figure 6B). The uniform yellow signals from the fibrillar networks confirmed the presence of both HSA and Tf inside these matured fibrillar assemblies. We envisaged that the matured fibrils of HSA act as templates for the adsorption/deposition of native Tf due to the exposed fibrillar surface of HSA within the heterogeneous droplet phase. To validate this argument, we performed a control experiment by mixing preformed HSA fibrils with the liquid-like droplets of Tf under similar experimental conditions (Figure 6E). CLSM measurements revealed time-dependent spontaneous adsorption of Tf droplets on the matured fibrils of HSA and upon contact, Tf droplets disintegrated to yield mixed fibrillar network with distinct yellow signals. Previously, Claessens and coworkers observed similar adsorption of negatively charged bovine serum albumin (BSA) on the positively charged amyloid fibrils of lysozyme and shown increased energy dissipation due to the spontaneous adsorption of protein.^64^ As expected, in the absence of any coalescence, surface immobilized heterogeneous droplets neither showed any nucleation nor any fibrillation upon maturation (Figure S19), further highlighting the importance of coalescence behind the aberrant aggregation.

Altogether, our findings not only highlight the critical role of coalescence-driven specific protein-protein interactions on the aberrant fibrillation of HSA within the homogeneous and heterogeneous droplet phase but also showcase the regulatory role of small ligands on the feasibility and kinetics of protein aggregation via modulating the soft protein-protein interactions (Scheme 2).

**Scheme 2.**
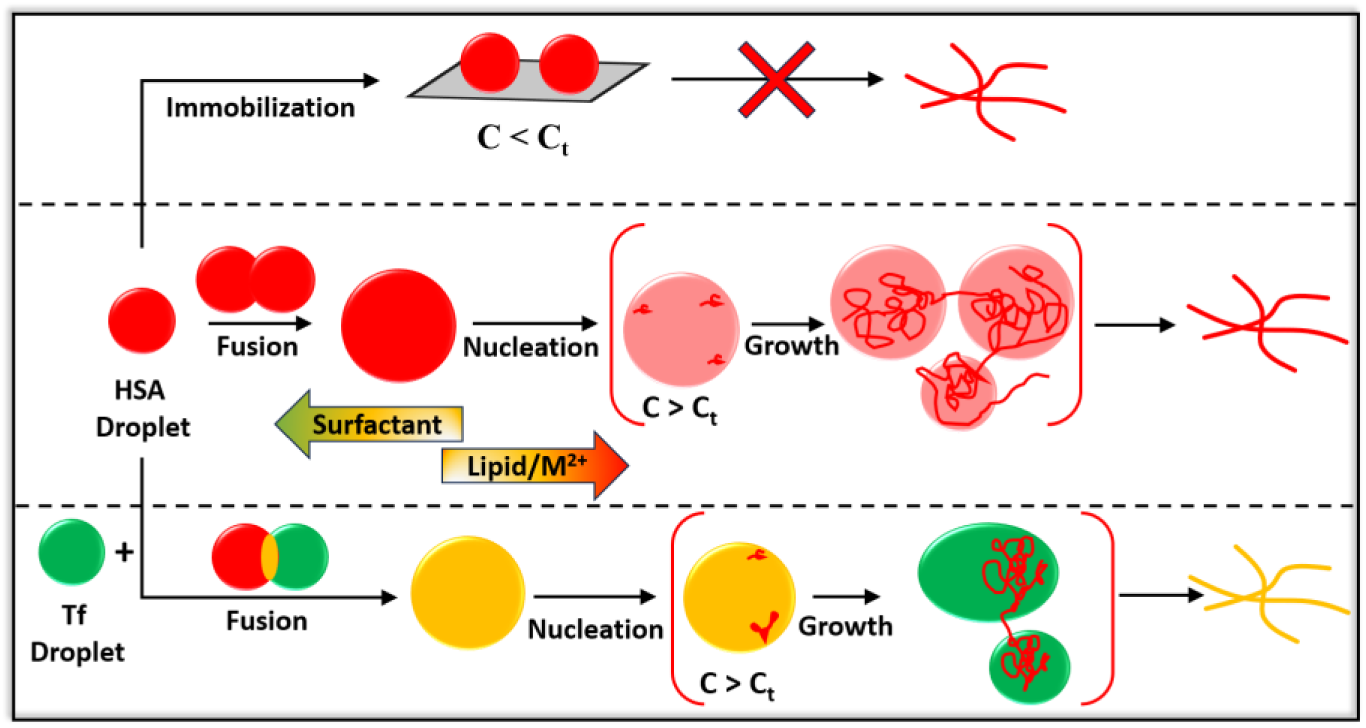
Schematic Illustration Showing Coalescence-Driven LSPT and its Modulation in a Homogeneous and Heterogeneous Droplet Assembly.

We show that spontaneous coalescence events help to cross a threshold protein concentration (C_t_) required for nucleation and subsequent growth. Importantly, surface immobilization or binding of charged surfactants prevents the coalescence events and subsequent LSPT of HSA. Further, our findings reveal that the early nucleation occurs mainly at the droplet surface and the nucleated structures undergo free diffusion within the boundaries of droplets. We discover that soft multivalent protein-protein interactions can efficiently dictate the organization of biomolecules within the multicomponent heterogeneous droplets. Additionally, our observation of spontaneous adsorption of functional biomolecules on the fibrillar surface within the heterogeneous droplet phase highlights the adverse effect of aberrant LSPT of biomolecules on the structure and function of other functional proteins in a heterogeneous crowded environment. Although, many efforts have been made in recent times to understand the role of heterotypic protein-protein interactions on the LLPS and LSPT in multicomponent droplets,^11,12,65^ the present study is unique to capture the microscopic transient intermediates and showcase the impact of specific homotypic protein-protein interactions on the spatial organization of the biomolecules and their aggregates within the multicomponent heterogeneous droplet assembly of two functional proteins.

### Summary

In the present study, we have investigated the detailed mechanistic aspects and microscopic events associated with the LSPT of functional protein assemblies involving metastable liquid-like droplets using a range of spectroscopic and microscopic techniques. The LSPT of HSA proceeds via formation of liquid-like droplets which subsequently transforms to amyloid-like fibrillar assembly via nucleation-dependent growth mechanism. Using CLSM imaging, we have shown that the initial nucleation originates at the droplet surface/interface and the subsequent elongation occurs exclusively inside the individual droplets. Further, we have demonstrated that spontaneous coalescence of droplets leads to the enhanced local protein concentrations inside the individual droplets and nucleation starts only after crossing a threshold protein concentration. These findings are authenticated using single droplet Raman measurements, which indicate gradual change in the local environment of the liquid-like droplets during the LSPT of HSA. In addition, we have illustrated the remarkable role of small ligands in regulating the feasibility and kinetics of LSPT of HSA due to the changes in its native conformation, which alter the intermolecular protein-protein interactions. Finally, to mimic the highly crowded and heterogeneous environment of living membraneless cellular organelles, we have designed a dynamic heterogeneous droplet assembly and shown heterogeneous nucleation and fibrillation of HSA which acts as template for the subsequent adsorption of structurally stable Tf to yield a mixed fibrillar assembly. Our present work thus opens up new avenues to explore and regulate the spatiotemporal behavior of metastable protein condensates on their aggregation pathways.

## ASSOCIATED CONTENT

### Supporting Information

Experimental Methods; turbidity assay; temporal evolution of CD spectra; temporal evolution of FTIR spectra of HSA; deconvoluted FTIR spectra with associated vibrational modes; ThT assays as a function of HSA concentration; fluorescence enhancement of ThT upon aging with HSA droplets; CLSM images of fusion, surface wetting, and dripping; CLSM image of HSA droplets after 5 min of mixing with 10% PEG; CLSM image of HSA droplet between d-5 and d-6; CLSM images during LSPT after removal of dilute supernatant phase; size distribution histogram of HSA droplets in the presence of 10 mM SDS and 2 mM CTAB upon aging; CLSM images of HSA droplets at lower concentration of SDS and CTAB; CLSM images during LSPT of HSA droplets in the presence of 10 mM Ca^2+^; size distribution histogram of HSA droplets in the presence of DPPC, Cu^2+^, and Ca^2+^; changes in the CD spectra of HSA in the presence of different ligands; secondary structure of HSA in the presence of different ligands; CLSM images of surface aged droplets upon aging; characteristic Raman vibrational modes of HSA; CLSM images of heterogeneous droplets of HSA and Tf upon surface aging; tabulated Raman peak positions of HSA (PDF)

## AUTHOR INFORMATION

### Corresponding Author

**Tushar Kanti Mukherjee** - Department of Chemistry, Indian Institute of Technology (IIT) Indore, Indore 453552, Madhya Pradesh, India;

### Author

**Chinmaya Kumar Patel** – Department of Chemistry, Indian Institute of Technology (IIT) Indore, Indore 453552, Madhya Pradesh, India

**Abhradip Mallik** – Department of Chemistry, Indian Institute of Technology (IIT) Indore, Indore 453552, Madhya Pradesh, India

**Deb Kumar Rath** – Department of Physics, Indian Institute of Technology (IIT) Indore, Indore 453552, Madhya Pradesh, India

**Rajesh Kumar** – Department of Physics, Indian Institute of Technology (IIT) Indore, Indore 453552, Madhya Pradesh, India

### Notes

The authors declare no competing financial interests.

## Acknowledgements

The authors acknowledge Indian Institute of Technology (IIT) Indore for providing financial support, and infrastructure. This work is financially supported by Council of Scientific and Industrial Research (CSIR) grant no. 01/3108/23/EMR-II. The authors acknowledge SIC, IIT Indore, for instrumental facilities. The Raman facility received from the Department of Science & Technology (DST), Government of India, under the FIST scheme (Grant no. SR/FST/PSI-225/2016) is gratefully acknowledged. C.K.P. and D.K.R. acknowledge Ministry of Education (MoE) and Prime Minister’s Research Fellowship (PMRF), India, respectively, for research fellowships.

## Notes

### Competing Interest Statement

The authors have declared no competing interest.

