## Supporting Information for "Coalescence-Driven Local Crowding Promotes Liquid-to-Solid-Like Phase Transition in a Homogeneous and Heterogeneous Droplet Assembly"

**TABLE OF CONTENTS**


| S.N. | | Page No. |
|---|---|---|
| 1 | Methods | S3-S6 |
| 2 | Supporting Figures | S7-S25 |
| 3 | Table S1 | S26 |
| 4 | References | S27 |

### Methods

#### 1. Preparation of Solutions

All the samples in this study were prepared in phosphate buffer saline. Phosphate-buffer saline (PBS, 50 mM, pH 7.4) was prepared using Milli-Q water. To this buffer solution, 50 mM NaCl and 0.02% sodium azide were added. Separately, a 10% (w/v) PEG solution was prepared by diluting a 40% (w/v) stock solution of PEG 8000 in PBS.

#### 2. Cleaning of Coverslips and Glass Slides for Microscopy

The coverslips and glass slides were first cleaned with 2% Hellmanex III followed by cleaning with chromic acid for 30 min. Each of these cleaning steps was followed by repeated washing with Milli-Q water. Finally, these washed coverslips and slides were rinsed with methanol and dried in vacuum oven.

#### 3. Labelling of Protein with Fluorescent Dyes

The concentration of HSA and Tf was estimated spectrophotometrically using the reported extinction coefficient of 35,700 $M^{-1}$ $cm^{-1}$ and $1.04 \times 10^5$ $M^{-1}$ $cm^{-1}$ respectively at $\lambda$ = 280 nm.[1,2] HSA was labeled with RBITC dye according to an earlier reported method.[3] In short, 3 mM HSA was mixed with RBITC in a molar ratio of 1:1 ([HSA]: [RBITC]). The mixture was incubated for 4 h at room temperature followed by 6 h at 4 °C on a magnetic stirrer with slow rotation. After the completion of the reaction, the unconjugated dyes were removed using dialysis (molecular weight cut-off 3.5 kDa) against 50 mM PBS at 4 °C for 12 h with regular buffer exchange in 2 h intervals. The estimated labeling efficiency was found to be ~77.3-84.1%.[4] Similar process was followed for labeling of Tf with FITC.

#### 4. Sample Preparation

**Solution Phase Aging**: HSA and Tf solutions were prepared in glass vials with phosphate buffer saline (PBS, pH 7.4) in the absence and presence of 10% PEG and incubated at 37 °C for a period of 14 days. For confocal imaging, the liquid samples were drop-cast onto a cleaned cover glass and sandwiched with a Blue Star coverslip. The edges of the coverslips were sealed with a minimal amount of commercially available nail paint.

**Surface-aging (surface immobilization)**: For surface aging experiments, 20 µL aliquot of the sample solution was drop-cast over a cleaned coverslip and kept inside the incubation chamber at 37 °C for a period of 1–14-days.

#### 5. LLPS Assay of HSA in the Presence of PEG 8000

LLPS at pH 7.4 was examined by equilibrated 500 µM HSA with 10% (w/v) PEG 8000 in PBS solutions at different time intervals. The samples were prepared in 5 mL glass vials and kept for incubation at 37 °C inside a constant temperature incubator. LLPS and LSPT assays were performed under CLSM.

#### 6. Turbidity Measurements

The turbidity measurements of the HSA solutions in the presence of 10% PEG 8000 were performed by recording the UV-vis absorption spectra. The turbidity of the equilibrated binary mixture was calculated using the following equation,

$$T = 100 - (100 \times 10^{-A}) \qquad (1)$$

where *T* is the turbidity and *A* is the absorbance at 350 nm.

#### 7. ThT Binding Assay

Stock solution of 2 mM ThT was prepared in pH 7.4 PBS. 20 µM ThT was used for the study of ThT binding assay. The ThT binding assay of HSA was carried out by Horiba FluoroMax Fluorometer using the excitation laser 450 and emission in the range of 460-550 nm. The working solutions were diluted by 20-fold before spectral measurements and maximum intensity at 480 nm was plotted against time.

The aggregation kinetics of HSA follows a sigmoidal curve, characteristic of amyloid fibril formation. This curve can be divided into three phases: the lag phase, the growth phase, and the saturation phase. The lag time was estimated as per the previously published protocol.[5] The lag time was estimated by extending the tangent at $t_m$ (inflection time,) to the baseline or time axis.

### 8. Ligand Binding Study

Stock solution of SDS, CTAB, DPPC lipid, $CuCl_2$, and $CaCl_2$ were prepared in pH 7.4 PBS. The samples were prepared by equilibrating 500 µM HSA in the presence of either 10 mM SDS, 2 mM CTAB, 1 mM DPPC lipid, 250 µM $CuCl_2$, or 10 mM $CaCl_2$ in PBS for 1 h at 37 °C, followed by the addition of 10% PEG and stored in 37 °C incubation chamber.

### 9. Student *t*-test Analysis

Statistical analyses were performed via a two-tailed, unpaired Student's *t*-test with ***, $P$ value $< 0.001$; **, $P$ value $< 0.01$, and not significant (NS), $P > 0.05$, using Excel software.

### 10. Estimation of Concentration Using Vibrational Raman Spectroscopy

The protein concentration inside the droplets was measured as described previously.[6] In brief, we estimated the protein concentration inside the individual droplets by Raman spectroscopy using the characteristic phenylalanine (Phe) peak at 1005 $cm^{-1}$ as standard which corresponds to the symmetric breathing of the benzyl ring of phenylalanine. Next, we applied this method to measure the concentration of HSA inside the condensed phase by focusing the laser beam on free HSA solution (500 µM HSA) and at the centre of the time-dependent HSA droplets (500 µM HSA in the presence of 10% PEG at d-1 and d-5). 532-nm NIR laser was used for excitation, with an exposure time of 10 s and 40 mW laser power was used in the Raman settings and recorded in between 850-1150 $cm^{-1}$.

### 11. Estimation of Protein Concentration Using Centrifugation

Droplet phase and dilute phase concentrations were measured using centrifugation based on a previously described method.[7] To estimate the droplet phase concentration, a solution of 500 µM HSA in the presence of 10% PEG was prepared. After incubating the sample at 37 °C for d-1 at constant stirring, it was centrifuged at ~18,000 rpm for 30 minutes. The supernatant was carefully removed in steps, and the droplet phase (~15 µL) was resuspended in a known volume of denaturation buffer (6M GdmCl, 20 mM sodium phosphate), with a 200-fold dilution. The absorbance at 280 nm was measured using an extinction coefficient of $\varepsilon_{280}$ = 35,700 $M^{-1}cm^{-1}$. The dilute phase concentration was determined by measuring the absorbance of the supernatant. Data were calculated from three independent experiments under the same conditions.

### 12. Formulation of Heterogenous Droplet Assembly

**Solution Phase Aging**. we mixed phase-separated droplets of 500 µM RBITC-labeled HSA and 50 µM FITC-labeled Tf in the presence of 10% PEG in a 1:1 ratio and equilibrated at 37 °C for a period of 1–12 days and aged in solution phase for CLSM monitoring.

**Surface Aging**. For surface aging experiments, a 20 µL aliquot of the heterogeneous droplet assembly (comprising droplets of 500 µM RBITC-labeled HSA and 50 µM FITC-labeled Tf) was drop-cast onto a cleaned glass slide. The slide was then placed in an incubation chamber at 37 °C for 14 days for subsequent study using CLSM.

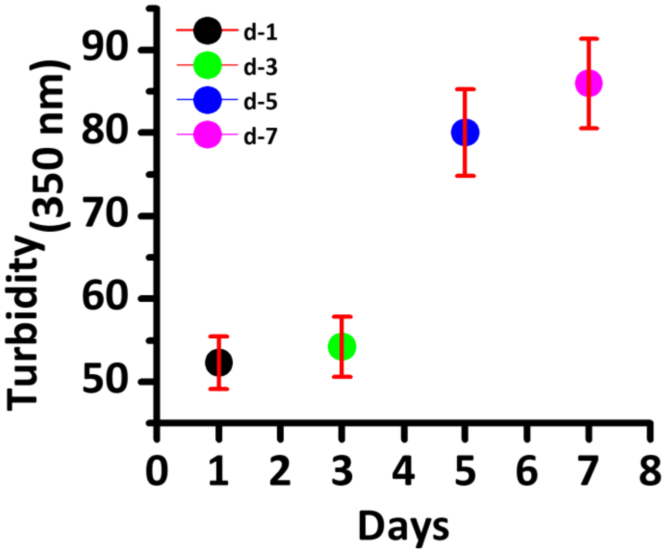


**Figure S1.** Changes in the turbidity of 500 µM HSA in the presence of 10% PEG measured at 350 nm as a function of aging. Data represent mean ± s.e.m. for three independent experiments (n = 3).

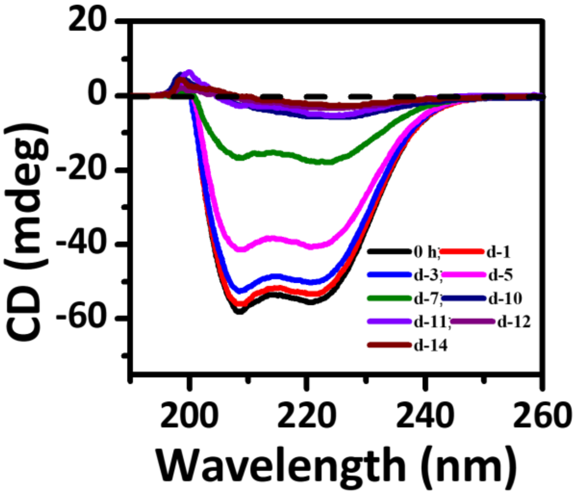


**Figure S2.** Changes in the CD spectra of 500 μM HSA in the presence of 10% PEG over a period of 14 days. The working solutions were diluted by 100-fold before the measurements.

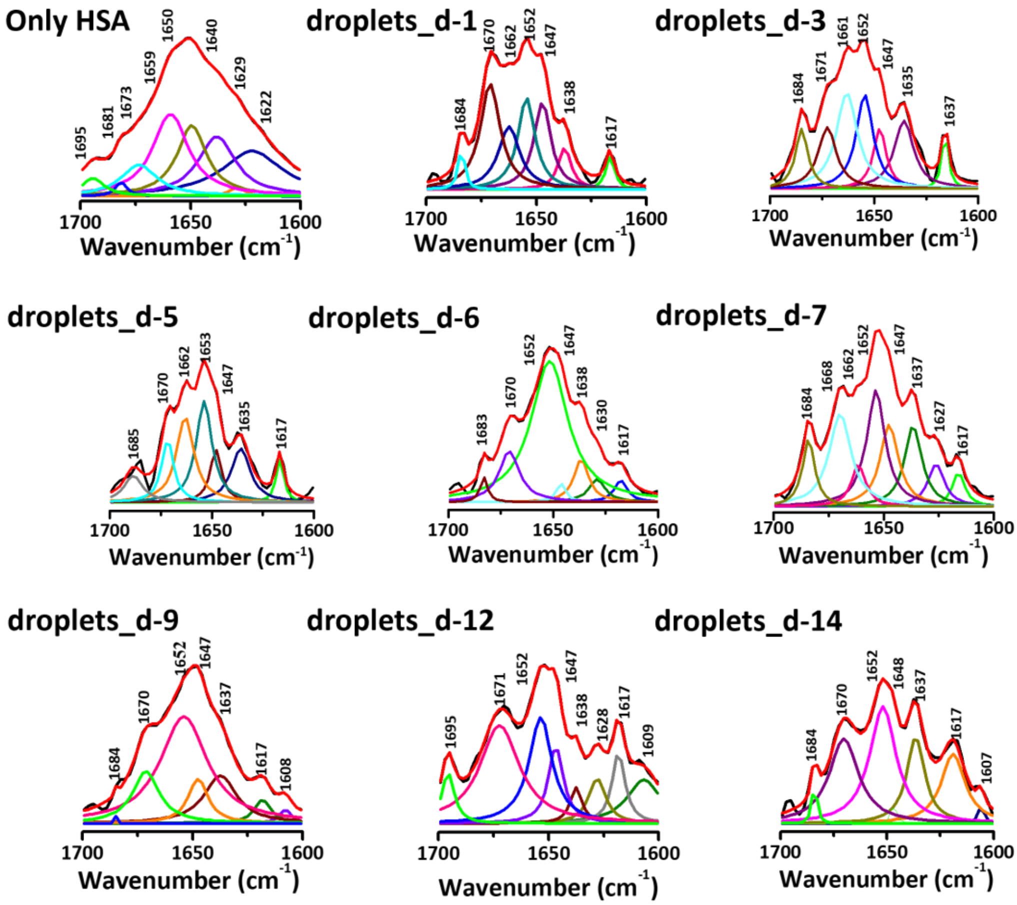


**Figure S3.** Changes in the deconvoluted FTIR spectra of 500 μM HSA in the absence and presence of 10% PEG over a period of 14 days.

(A)

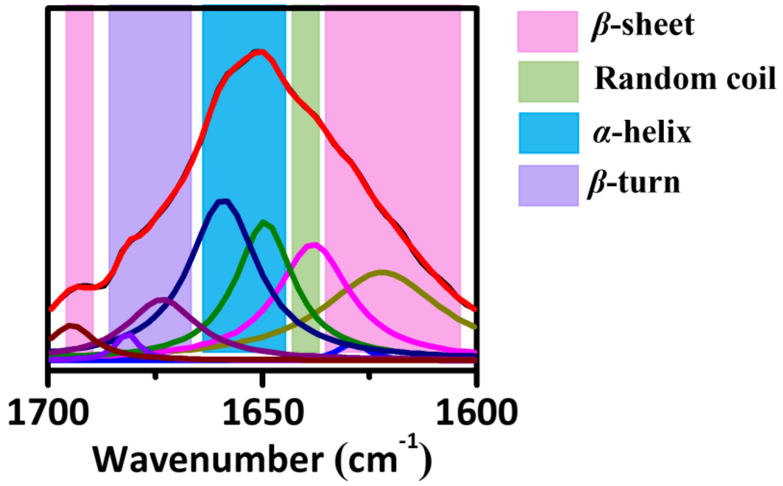


(B)

| Wavenumber ($cm^{-1}$) | Secondary structure assignment |
|---|---|
| 1610-1639; 1691-1696 | $\beta$-sheet |
| 1641-1648 | Random coil |
| 1649-1664 | $\alpha$-helix |
| 1668-1686 | $\beta$-turn |

**Figure S4.** (A) Deconvoluted FTIR spectrum and (B) associated secondary structures of HSA in the amide I band region. Secondary structure contents were estimated by calculating the relative area under each peak using origin 8.1 software.

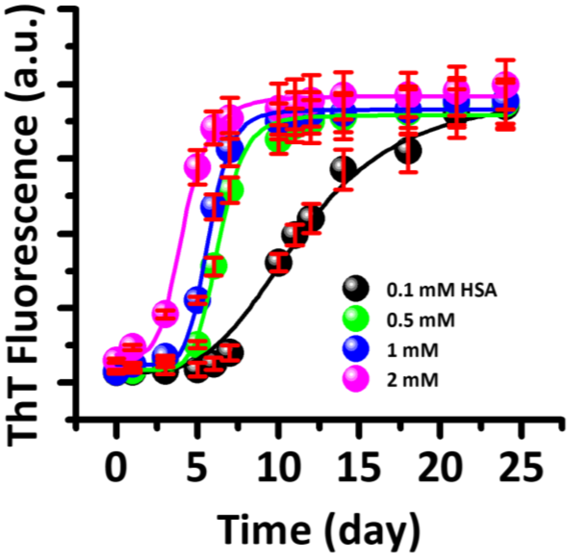


**Figure S5.** Aggregation kinetics monitored using ThT fluorescence ($\lambda_{ex}$ = 450 nm; $\lambda_{em}$ = 480 nm) as a function of HSA concentrations in the presence of 10% PEG. The working solutions were diluted by 20-fold before the spectral measurements. Data represent mean ± s.e.m. for three independent experiments (n = 3).

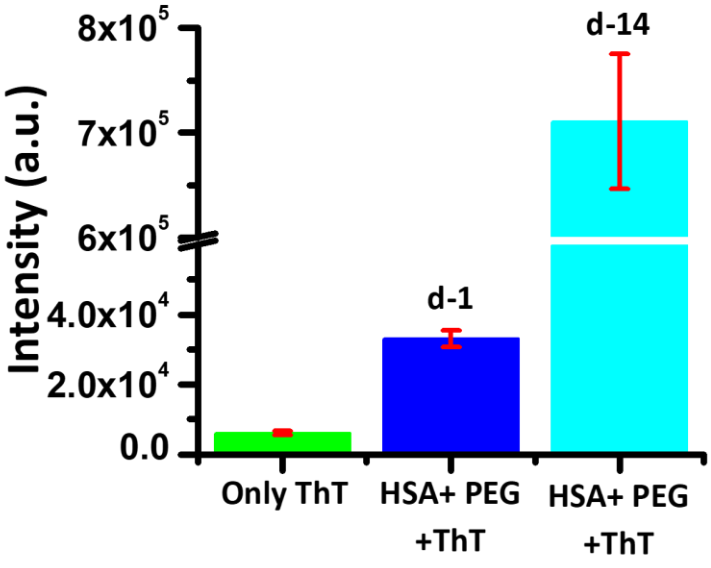


**Figure S6.** Changes in the ThT fluorescence intensity ($\lambda_{ex}$ = 450 nm; $\lambda_{em}$ = 480 nm) in the absence and presence of HSA droplets (500 µM HSA in the presence of 10% PEG) at d-1 and d-14. The working solutions were diluted by 20-fold before the spectral measurements. Data represent mean ± s.e.m. for three independent experiments (n = 3).

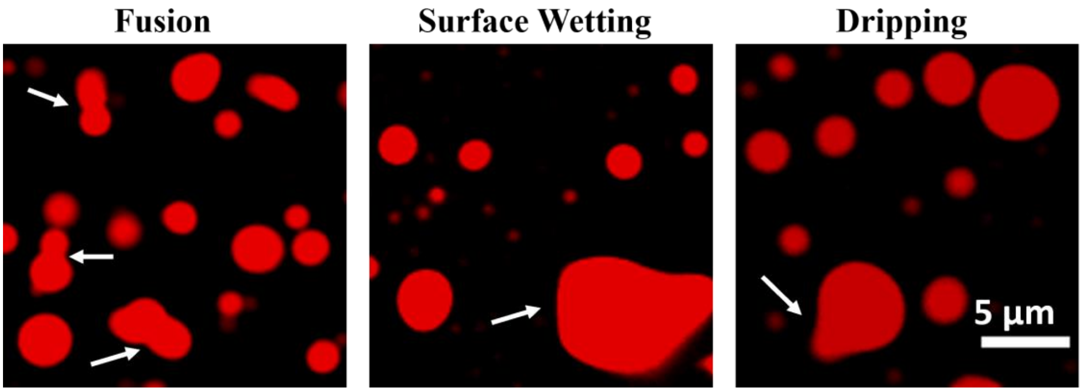


**Figure S7.** CLSM images showing the fusion, surface wetting, and dripping behaviours of RBITC-labeled HSA droplets.

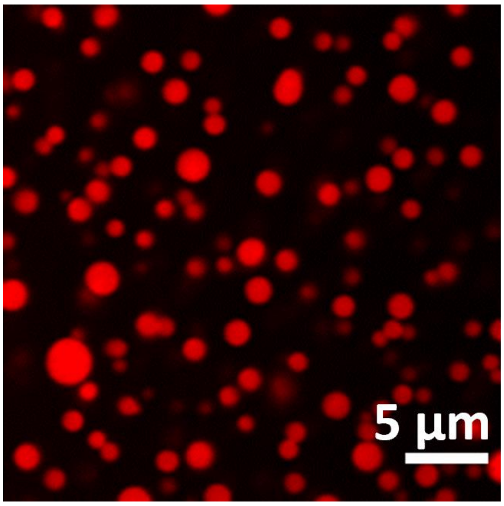


**Figure S8.** CLSM image of RBITC-labeled HSA droplets (500 µM HSA in the presence of 10% PEG) formed within 5 min of mixing.

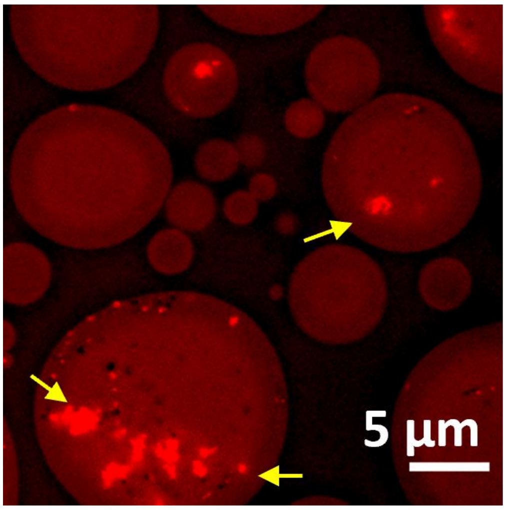


**Figure S9.** CLSM image showing the presence of aggregates (nuclei and clusters) within the RBITC-labeled HSA droplets upon aging between d-5 and d-6. Yellow arrows indicate the aggregates inside the droplets.

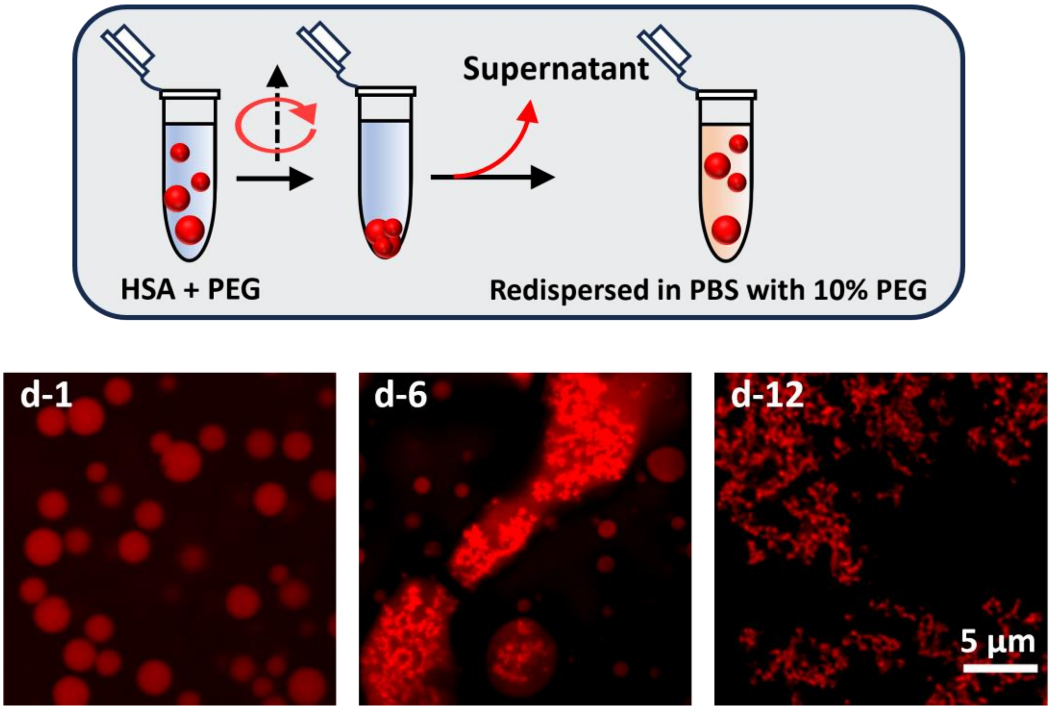


**Figure S10.** Confocal images showing the LSPT of RBITC-labeled HSA droplets (500 µM HSA in the presence of 10% PEG) prepared after replacing the dilute supernatant phase with equal volume of PBS and 10% PEG.

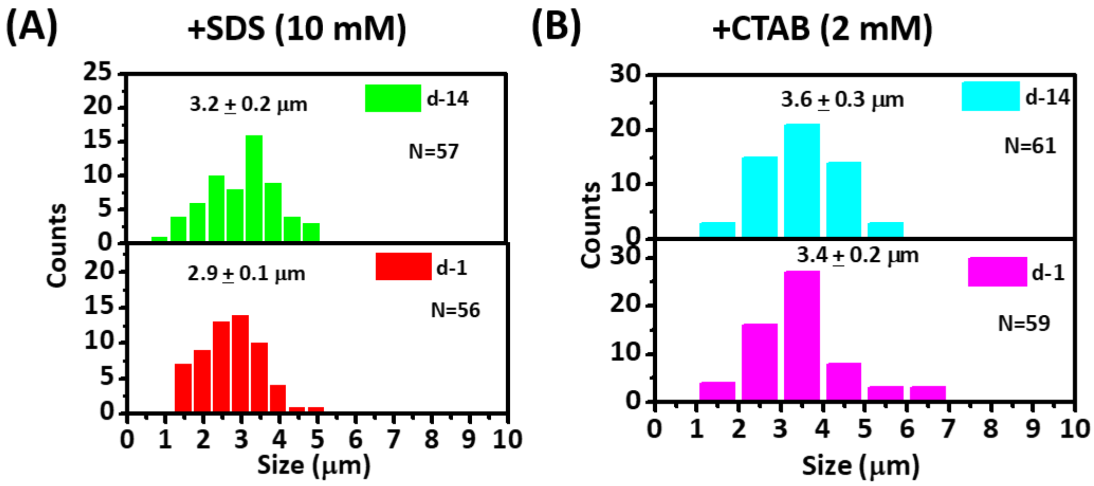


**Figure S11.** Size distribution histograms of HSA droplets in the presence of (A) 10 mM SDS, and (B) 2 mM CTAB estimated from CLSM images at d-1 and d-14. Data represent mean ± s.e.m. for three independent experiments (n = 3).

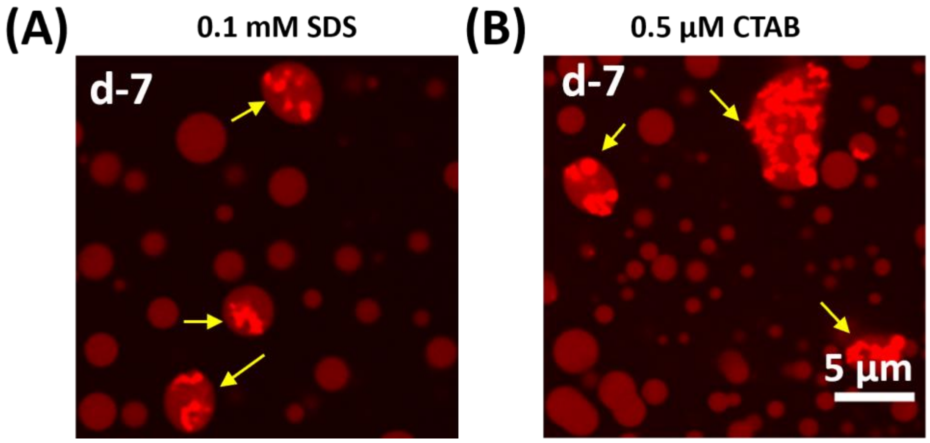


**Figure S12.** CLSM images showing the aggregation of HSA within the RBITC-labeled HSA droplets in the presence of (A) 0.1 mM SDS, and (B) 0.5 μM CTAB at d-7. Yellow arrows indicate the presence of aggregates within the droplets.

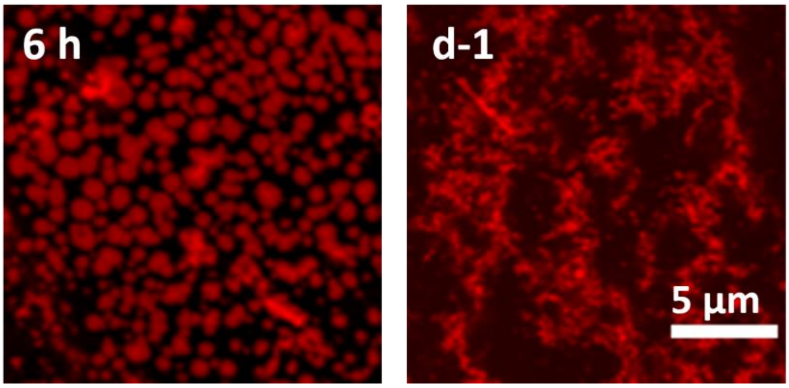


**Figure S13.** CLSM images of RBITC-labeled HSA droplets in the presence of 10 mM $Ca^{2+}$ upon aging.

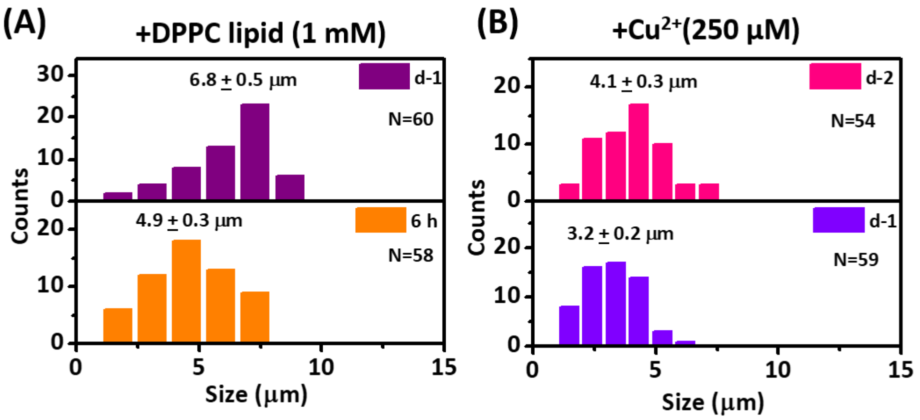


**Figure S14.** Size distribution histograms of HSA droplets in the presence of (A) 1 mM DPPC, and (B) 250 µM $Cu^{2+}$ estimated from CLSM images at different time intervals. Data represent mean ± s.e.m. for three independent experiments (n = 3).

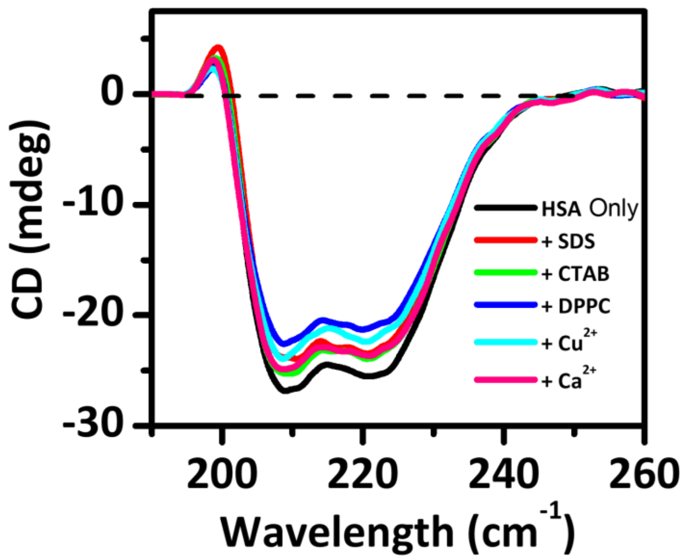


**Figure S15.** Changes in the CD spectra of HSA in the presence of 10 mM SDS, 2 mM CTAB, 1 mM DPPC, 250 μM $Cu^{2+}$, and 10 mM $Ca^{2+}$.

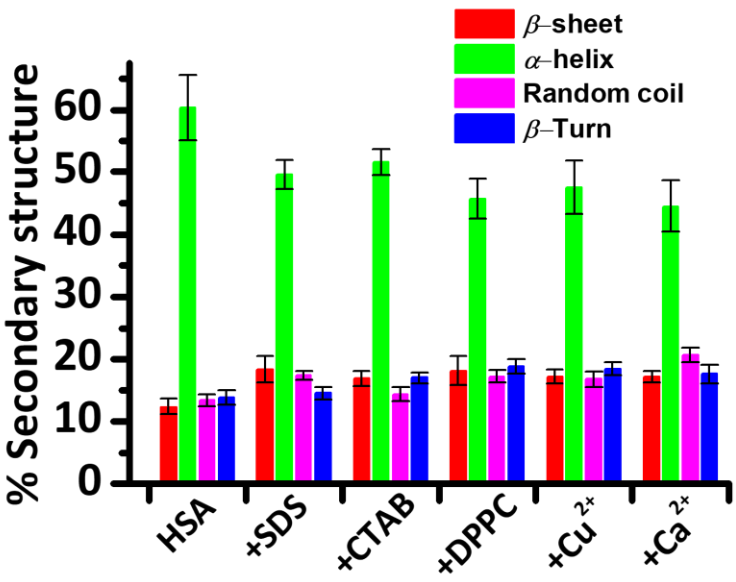


**Figure S16.** Secondary structure contents of HSA in the presence of 10 mM SDS, 2 mM CTAB, 1 mM DPPC, 250 μM $Cu^{2+}$, and 10 mM $Ca^{2+}$ estimated from deconvoluted FTIR spectra. Data represent mean ± s.e.m. for three independent experiments (n = 3).

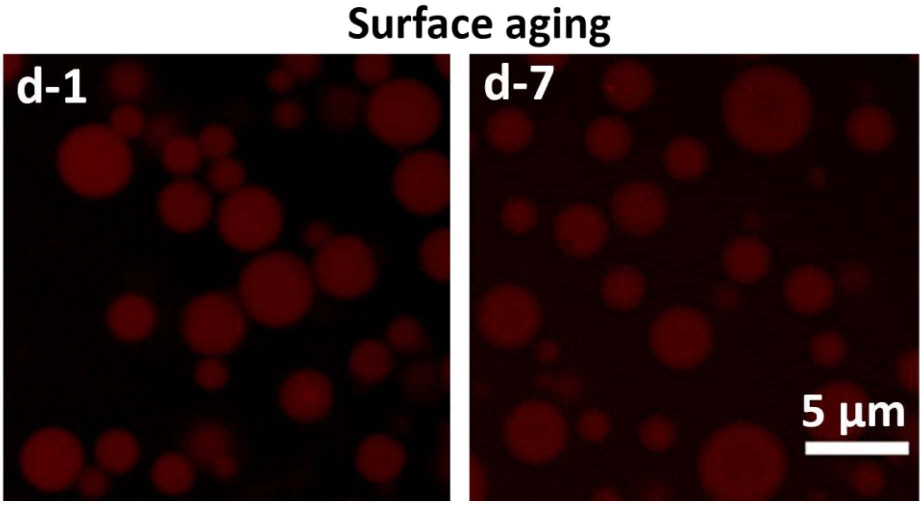


**Figure S17.** CLSM images of surface-aged HSA droplets (500 µM HSA in the presence of 10% PEG) at d-1 and d-7.

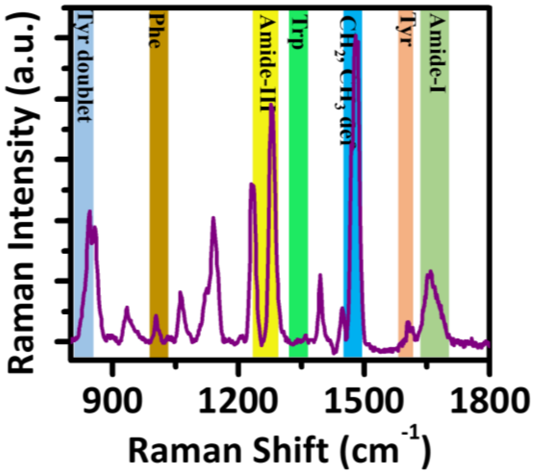


**Figure S18.** Representative vibrational Raman spectrum of HSA with associated vibrational features.

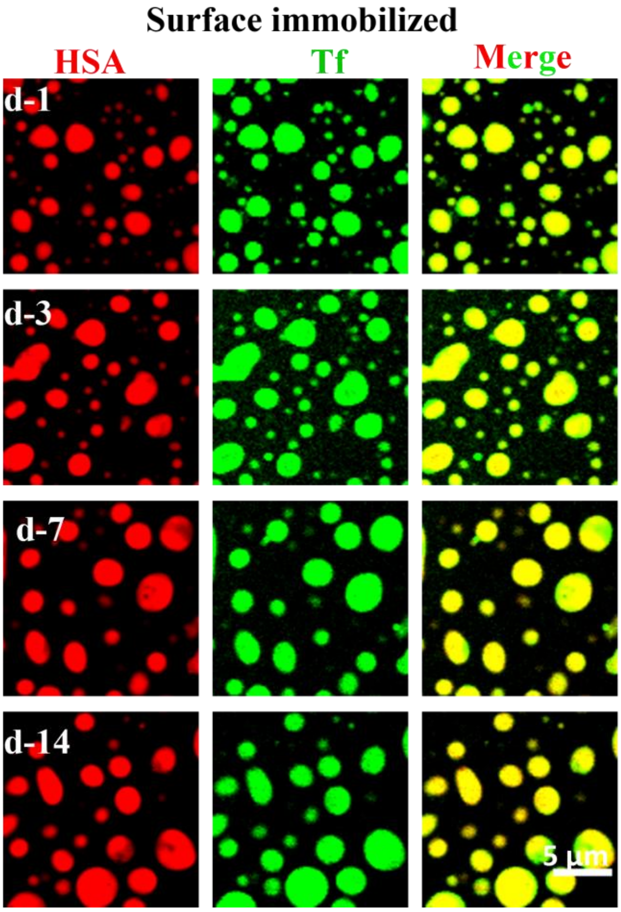


**Figure S19.** CLSM images of heterogenous droplets containing RBITC-labeled HSA and FITC-labeled Tf upon surface aging.

**Table S1. Vibrational Raman bands and associated vibrational modes of HSA**

| **Peak position assignment** | **band** |
|---|---|
| **Amide I** (primarily due to C=O stretching mode) | 1630-1700 $cm^{-1}$ |
| **Amide III** (due to in-plane N-H bending and C-N stretching motions) | 1220-1320 $cm^{-1}$ |
| **Phenylalanine** (due to ring breathing vibrations) | 1000-1008 $cm^{-1}$ |
| **Tryptophan** | 1338 $cm^{-1}$<br>(Fermi doublet at 1360 $cm^{-1}$ and 1340 $cm^{-1}$) |
| **Tyrosine**<br>**a. Fermi doublet at 850 $cm^{-1}$ and 830 $cm^{-1}$**<br>**b. Ring stretching mode** | <br>$I_{850}/I_{830}$ is an indicator of solvent mediated hydrogen bonding propensity of the phenolic (-OH) group<br>1600 $cm^{-1}$ and 1617 $cm^{-1}$ |
| **Backbone $CH_2/CH_3$ deformations** | 1440-1470 $cm^{-1}$ |
